# Malaria drives unique regulatory responses across multiple immune cell subsets

**DOI:** 10.1101/2022.11.16.516822

**Authors:** Nicholas L. Dooley, Tinashe G. Chabikwa, Zuleima Pava, Jessica R. Loughland, Julianne Hamelink, Kiana Berry, Dean Andrew, Megan S.F. Soon, Arya SheelaNair, Kim A. Piera, Timothy William, Bridget E. Barber, Matthew J. Grigg, Christian R. Engwerda, J. Alejandro López, Nicholas M. Anstey, Michelle J. Boyle

## Abstract

*Plasmodium falciparum* malaria results in immunoregulatory responses across multiple cell subsets, which protects the individual from inflammatory mediated immunopathogenesis. However, these anti-inflammatory responses also hamper the development of effective anti-parasitic immunity. Understanding malaria induced tolerogenic responses in specific cell subsets may inform the development of strategies to boost protective immunity during drug treatment and vaccination. Here, we analysed the immune landscape with single cell RNA sequencing of peripheral blood mononuclear cells during falciparum malaria and at convalescence in children and adults from a low malaria transmission area in Malaysia. To understand malaria driven changes specific to each immune cell subset, we interrogated transcriptional changes in sub-clustered major immune cell types during infection. We found that malaria drove development of immunosuppressive monocytes, alongside NK and γδ T cells which regulated inflammatory function but maintained cytolytic capacity. IL10-producing CD4 T cells and IL10-producing regulatory B cells were also induced. Type I interferon responses were identified across all cell types, linking Type I interferon signalling with the induction of immunoregulatory networks during malaria. Together, these findings provide insights into cell-specific and shared immunoregulatory changes induced during malaria, and provides a data set resource for additional analysis of anti-parasitic immunity and disease pathogenesis.

## INTRODUCTION

*Plasmodium falciparum* causes significant disease burden globally, with >240 million cases of malaria and >600 000 deaths reported in 2020 (*1*). In areas of high malaria transmission, anti-disease or tolerogenic immunity develops relatively rapidly, with children rarely experiencing recurrence of severe malaria (*2*). However, anti-parasitic immunity and protection from mild diseases develops more slowly, with children experiencing multiple patent infections throughout childhood before developing levels of protection that control parasite growth to sub-patent levels. These phenomena are thought to be linked, with the slow acquisition of anti-parasitic immunity attributed to tolerogenic responses across multiple cell subsets required for robust adaptive immune development. These tolerogenic mechanisms may also contribute to reduced malaria vaccine efficacy in exposed populations. As such, a better understanding of tolerogenic immune responses during infection may inform the development of more effective vaccine strategies for areas of high malaria endemicity.

Malaria driven tolerogenic and immunoregulatory responses allow parasite persistence by evading and disrupting anti-parasitic mechanisms employed by both innate and adaptive immune cells. Multiple studies have shown that monocytes and dendritic cells (DCs), that are initiators of the immune responses, are tolerized during malaria. During experimental human malaria and natural infection, monocytes and DCs have reduced responsiveness to toll-like receptor (TLR) stimulation and antigen presentation is suppressed (*3–6*). Increased IL-10 production (*7*) and higher frequencies of monocytes with a regulatory phenotype are also detected in children and adults from malaria endemic areas (*8, 9*). Tolerogenic phenotypes have also been reported in other innate cells. For example, natural killer (NK) cells expressing the regulatory marker PD1, are expanded in populations in malaria endemic areas (*10*). Additionally, gamma-delta (γδ) T cells, particularly Vδ2^+^ subsets which are important innate inflammatory responders to malaria parasites, become tolerized in children in high transmission settings (*11, 12*). Immunoregulation also exists in adaptive cell responses to malaria. Within the CD4 T cell compartment, type 1 regulatory (Tr1) cells that co-produce IFNγ and IL10 during malaria, dominate antigen-specific CD4 T cell responses in children in high endemic areas (*13–15*). These Tr1 cells develop rapidly during a primary malaria infection in previously naive adults (*16*). CD4 T cells also upregulate a number of additional inhibitory pathways during malaria infection, including expression of co-inhibitory receptors and production of TGFβ (*17–19*). Within the B cell compartment, multiple studies have shown that ‘atypical’ memory B cells expand in response to malaria (*20–22*). These cells have reduced functional capacity compared to ‘typical’ memory B cells (*23, 24*), and an immunoregulatory role for these cells in malaria is also possible. Malaria responsive regulatory B cells (Bregs), which produce IL10, have been reported in mouse models(*25*), but have not been identified in human malaria. While the drivers of tolerogenic cell responses are incompletely understood, Type I IFN signalling is key to the emergence of Tr1 CD4 T cells during malaria (*16*), and is recognised as both an activating and regulatory driver of the malaria immune response (*26*).

Transcriptional changes associated with immune cell tolerance have been reported in limited studies. For example, monocytes from Malian children and adults following parasite stimulation *in vitro* had reduced induction of *NFKB1* (positive regulator of inflammation) in tolerized adult cells (*8*), and transcriptional analysis of Vδ2^+^ γδ T cells in Ugandan children identified upregulation of multiple immunoregulatory pathways in highly exposed children (*11*). Additionally, a large whole-blood transcriptomic study revealed the upregulation of interferon responses, and that p53 activation in monocytes attenuated *Plasmodium*-induced inflammation and predicted protection from fever (*27*). However, to date, no studies have comprehensively investigated transcriptional signatures of malaria-driven tolerance across all cell subsets in the same individuals during infection.

The advent of single-cell RNA sequencing (scRNAseq) has allowed comprehensive analysis of distinct immune cell subsets during human infection, and identification of key changes driven by infection. For example, in HIV, scRNAseq revealed previously under-appreciated differentiation of proinflammatory T cells, prolonged monocyte major histocompatibility complex II (MHC II) upregulation, and NK cell cytolytic killing (*28*). Additionally, throughout the SARS-CoV-2 pandemic, rapid application of scRNAseq platforms provided important comprehensive understanding of cell type specific responses to both mild and severe infections, as well as similarities and differences to other diseases (*29–31*). To date, while scRNAseq has been applied to the malaria parasites (*32–34*), no comprehensive scRNAseq mapping of the immune landscape during malaria infection has been undertaken. In the present study, we applied scRNAseq to peripheral blood mononuclear cells (PBMCs) from patients during acute falciparum malaria and post treatment. Tolerogenic responses during acute infection were identified across multiple immune cell subsets, with key transcriptional changes confirmed at the protein level in additional patients. Together this study advances our understanding of the regulatory immune landscape during malaria and provides opportunities to manipulate these pathways for clinical advantage.

## RESULTS

### Altered immune cell profiles during acute malaria infection

To undertake a global analysis of the immune response during malaria infection, we performed droplet-based scRNAseq on peripheral blood mononuclear cells (PBMCs) from 6 individuals (age 6-24 years) with uncomplicated *P. falciparum* malaria at hospital presentation (day 0) and at 7 and 28 days after drug treatment, along with 2 healthy adult endemic controls (ages 20 and 27 years) (**Fig. 1A**, **Table S1**). We sequenced a total of 115 526 cells, with 106 076 cells passing quality control (QC; minimum of 220 genes expressed and <20% mitochondrial reads per cell). Due to a 10X Chromium wetting error, no quality cells were retained from one individual at the acute infection time point (ID child 1). Data were integrated to harmonize data sets across batch, donor and infection timepoints, and cell clusters visualized with uniform manifold approximation and projection (UMAP). Expression of canonical and lineage marker genes were used to annotate cell clusters into 15 high level cell states: CD14^+^ classical monocytes, CD16^+^ non-classical monocytes, classical dendritic cells (cDCs), plasmacytoid dendritic cells (pDCs), CD4 T cells, CD8 T cells, γδ T cells, NKT cells, B cells, plasma cells, proliferating cells [which appeared to be of mixed cell types], hematopoietic stem and progenitor cells (HSPCs), platelets, and one unidentified cluster (**Fig. 1B/C, Table S2)**. The relative proportions of these cell clusters correlated strongly with the proportions of cells identified by flow cytometry analysis of the same cell samples (R=0.95, *p*<0.001, **Fig. 1D, Fig. S1**). During malaria infection, there were marked changes to the distribution of cell types, with relative increases in CD4 T cells and proliferating cells, and marked decreases in NK cells and γδ T cells (**Fig. 1E**). To characterize gene expression profiles during malaria infection, we performed differential gene expression analysis within each cell subset between acute infection (day 0), 7 and 28 days after treatment. (**Fig. 1F**). We observed the largest number of differentially expressed genes (DEGs) when comparing between day 0 and day 28, with monocytes and classical dendritic cells (cDCs) exhibiting the highest transcriptional changes relative to other cell types (**Fig. 1F, Tables S3A-C**). Large numbers of DEGs were also detected between day 0 and day 7, with a large proportion of these also detected 28-days post-treatment for each subset (for example, majority of DEGs for day 0 compared to day 7 and day 0 compared to day 28, were shared for CD14^+^ classical monocytes [64%], CD16^+^ non-classical monocytes [60%], cDCs [53%] and pDCs [47%]). As such, we focused subsequent analysis on DEGs identified between day 0 (acute malaria) and day 28 post treatment.

**Fig. 1.**
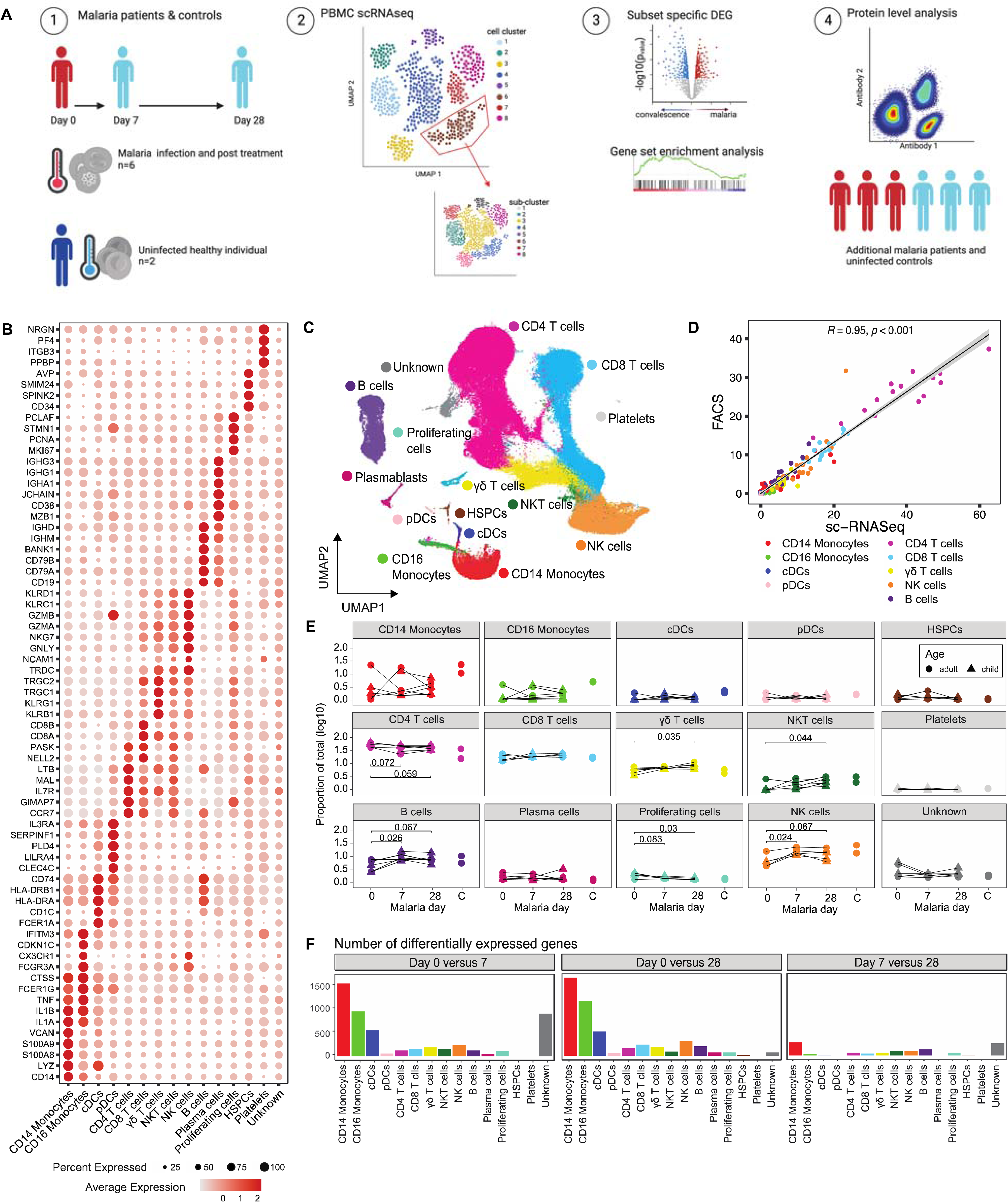
Single cell transcriptional landscape of malaria infection. (**A**) A schematic outline depicting workflow for sample collection and scRNAseq analysis. PBMCs were collected from falciparum malaria patients at day 0, and at day 7 and 28 post treatment (*n* = 6) and from malaria uninfected healthy controls (*n* = 2). Live PBMCs were analysed by 3’ 10X Chromium single cell sequencing, and cell types identified. For each cell type and sub-cluster, genes with differential expression between days were identified and analysed. Key findings were confirmed at the protein level in additional patients. (**B**) Dot plot of the mean expression of marker genes used to annotate cell types. (**C**) UMAP of all cells in integrated analysis. Cells are coloured by cell subtypes. (**D**) Correlation between relative proportion of cells identified by scRNAseq and flow cytometry analysis. Pearson’s R and p is indicated. (**E**) Relative proportion of identified subsets from scRNAseq analysis at day 0 during malaria, and day 7, and 28 days post-treatment, and in healthy uninfected control individuals. P-value is calculated by Mann Whitney U test between day 0 and subsequent time points. (**F**) Number of DEGs for each cell type between day 0/7, day 0/28 and day 7/28. See also Fig. S1 and Tables S1 to S3.

### Shared and subset-specific immunosuppressive signatures in monocytes and cDCs during malaria

We first analysed transcriptional changes to innate myeloid cells from day 0 to day 28 and identified 1674, 1182, 521 DEGs in CD14 classical monocytes, CD16 non-classical monocytes and cDCs, respectively (**Table S3B**). The high transcriptional activity of monocytes and cDCs during malaria was in contrast to the low number of DEGs detected in pDCs (*n*=60). Of pDC DEGs, 23% were associated with the long non-coding RNA family and 15% were histone genes (Table S3B). Low transcriptional activation of pDCs is consistent with our previous bulk-RNA sequencing analysis of isolated pDCs during experimental malaria infection, which showed that pDCs were transcriptionally stable during infection (*35*). Transcriptional changes during malaria in both monocyte subsets and cDCs were both shared and cell type specific (**Fig. 2A**). Both CD14^+^ and CD16^+^ monocytes were activated, with marked upregulation of innate cell activation genes such as *TOLL-LIKE RECEPTORS* (*TLRs*), *DISINTEGRIN and METALLOPEPTIDASE DOMAIN* (*ADAM9* and *ADAM10*) proteins(*36*), as well as alarmins, *S100A8* and *S100A9*, which are typically upregulated in monocytes under inflammatory conditions (*37*). In contrast, MHC class II HLA-DR genes were down-regulated in both CD14 and CD16 monocytes (**Fig. 2B,** Table S3B). High expression of S100A8/A9, along with *RETN*, *ALOX5AP* and reduced expression of MHC class II HLA-DR genes has recently been used to define immunosuppressive MS1 monocytes in sepsis (*38*) and proportional increases in these immune-suppressive monocytes has been detected in both sepsis and COVID-19 (*30*). Consistent with an enrichment of immunosuppressive monocytes during malaria infection, a number of inflammatory cytokines including *TNF, IL1α, IL1β, IL6, IL18* were markedly reduced at day 0, compared with 28 days post treatment (**Fig. 2B, Table S3B**). There was also reduced expression of multiple chemokine genes including *CCL2* (encoding MCP-1), *CCL3, CCL4, CCL5, CCL7, CXCL8* (encoding IL8), *CXCL2, CXCL3, CCL20, CXCL1* and *CXCL16* in both monocyte subsets, with a greater magnitude of reduction in CD14 monocytes. Consistent with reduced expression of these cytokine and chemokine genes, we also observed reduced expression of NK-κB family members, *NF-κB1*, *NF-κB2*, and *REL*, central transcriptional factors for pro-inflammatory gene induction (*39*). Immune-suppressive phenotypes were also detected in cDCs during infection, with notable down-regulation of HLA-DR genes *HLA-DRA and HLA-DRB1* along with multiple paralogues (*HLA-DPA1/B1* and *HLA-DQA1/B1*) in cDCs but not cytokine/chemokine genes. The reduction of HLA-DR genes in cDCs is consistent with previous reports showing reduced HLA-DR expression at the protein level on DCs during experimental (*3*), and naturally acquired malaria (*5, 40, 41*).

**Fig. 2.**
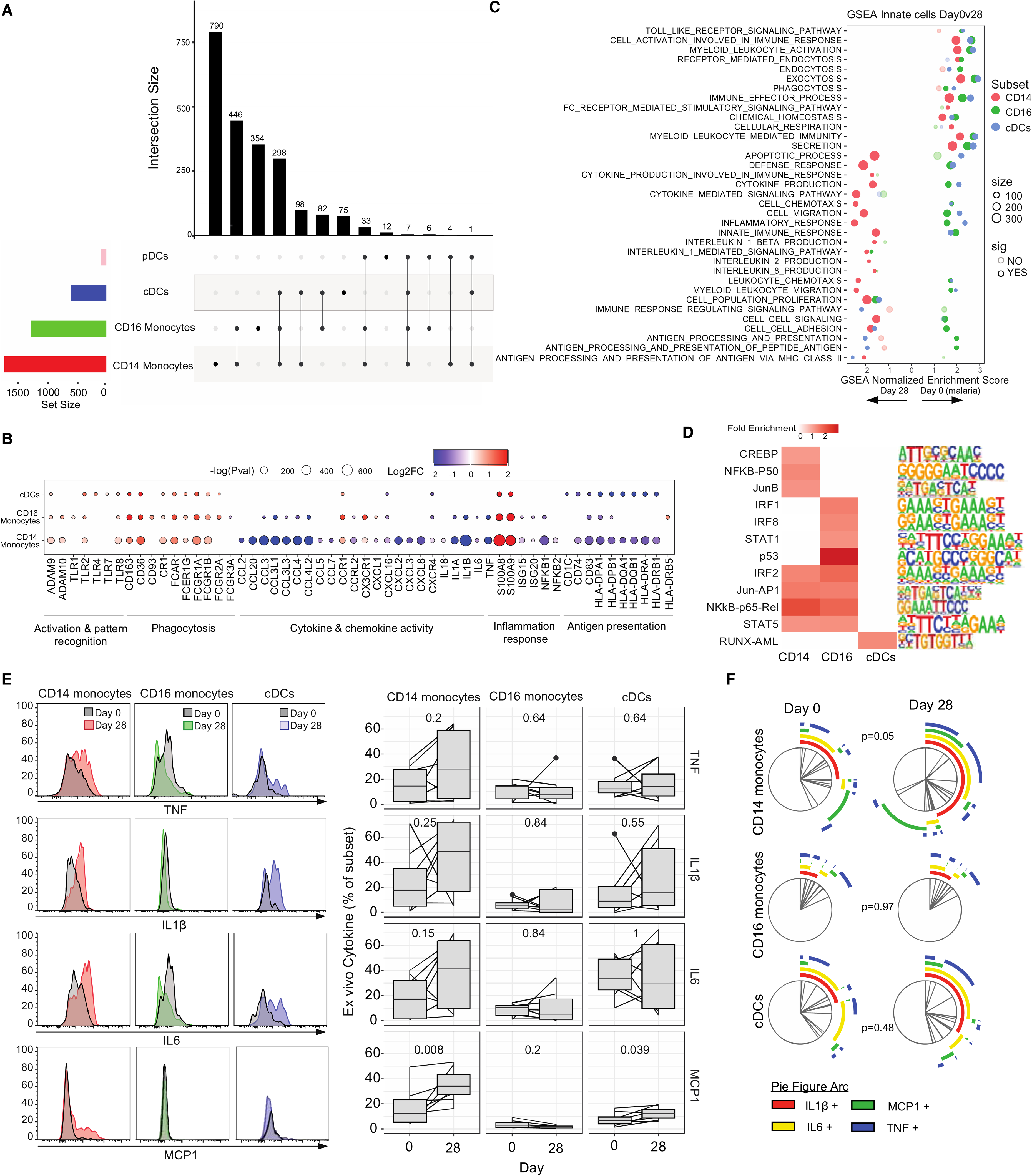
Immuno-suppressive signatures of innate cells during malaria. DEGs in CD14 and CD16 monocytes, and cDCs and pDCs were at day 0 compared to day 28 were identified. (**A**) Upset plot of shared and subset specific DEGs of each subset identified between day 0 and day 28. (**B**) DEGs of interest in monocytes and cDCs. Genes with known monocyte function are indicated. (**C**) GSEA of DEGs in CD14 and CD16 monocytes and cDCs. (**D**) Common and unique upstream regulators of DEGs in CD14 and CD16 monocytes and cDCs. (**E**) *Ex vivo* secretion of TNF, IL1β, IL6 and MCP1 was measured in CD14 and CD16 monocytes, and cDCs. Left panel – representative cytokine expression of a single individual at day 0 compared to day 28 in each subset. Right panel – population level expression of cytokine expression (*n*=8 day 0, *n*=8 day 28). P-value indicated is calculated by Wilcoxon signed rank test. (**F**) Co-expression of cytokines in each subset was analysed by SPICE. Expression graphed as Pie-figures. P-value indicated is calculated by Permutation test. See also Fig. S2 and Table S3.

In contrast to down-regulation of inflammatory gene signatures, multiple genes involved in pathogen recognition, scavenging and phagocytosis were upregulated in both monocyte subsets and cDCs during acute infection, including *CD163* and *FCGR1A* (encoding CD64/FcγRI) which we have previously shown to be upregulated at the protein level on both classical and non-classical monocytes during malaria (*4*) (**Fig. 2B**). Similarly, *FCAR* (encoding the Fc fragment of IgA receptor) was also upregulated on both monocyte subsets and cDCs, consistent with a possible role of IgA targeting antibodies in immunity against malaria (*42–44*). *CD93* and *FCGR2A* (encoding CD32/FcγRIIa) which mediate the enhancement of phagocytosis in monocytes and macrophages were upregulated during infection on CD16 monocytes and cDCs, but not CD14 monocytes. *COMPLEMENT RECEPTOR 1* (*CR1*), a membrane immune adherence receptor that plays a critical role in the capture and clearance of complement-opsonized pathogens including by erythrocytes and monocytes/macrophages, was upregulated on both CD14^+^ and CD16^+^ monocytes and cDCs during malaria. This transcriptional upregulation of *CR1* is in contrast to previous reports of down regulated CR1 on splenic monocytes/macrophages in murine models of malaria, and on CD16^+^ monocytes/macrophages in *P. falciparum* and *P. vivax* malaria patients from Peru(*45*). CD36, involved in antibody independent phagocytosis was also upregulated on both subsets and cDCs.

To further investigate shared and cell type specific transcriptional changes, DEGs were analysed using the Gene Set Enrichment Analysis (GSEA) and overrepresented upstream regulators identified. These analyses revealed both shared and cell specific gene signature enrichment and regulators (**Fig. 2C-D**). Consistent with DEGs for each subset, both monocyte subsets and cDCs were enriched for pathways such as secretion, phagocytosis, myeloid leukocyte mediated immunity and immune effector processes during malaria. In contrast, multiple subset specific pathways were also identified. For example, leukocyte migration and chemotaxis, inflammatory and defence response, and cytokine production were enriched at day 0 during malaria in CD16 monocytes and cDCs, but enriched at day 28 in CD14 monocytes. In contrast, pathways associated with antigen presentation were consistently enriched only in CD16^+^ monocytes during malaria. In agreement with cell type specific pathway enrichment, analysis of upstream regulators identified motifs that were shared between CD14^+^ and CD16^+^ monocytes including NFκB-p5, IRF2, STAT5 and Jun-AP1-binding *cis*-elements, but also regulators that were specific for each subset. For CD16 monocytes, this included p53, STAT1, IRF1, and IRF8-binding *cis*-elements. p53 was previously identified in bulk transcriptional analysis as an important contributor of innate cell responses and immunity to malaria (*27*). The enrichment of IFR1/8/2 and STAT1/5 are consistent with a key role of Type I IFN pathways in both activation and regulation of innate immune cell responses in malaria(*26*).

To assess some of these key findings at the protein level, *ex vivo* secretion of cytokine/chemokines TNF, IL1β, IL6 and MCP1 (*CCL2*) from CD14^+^ monocytes, CD16^+^ monocytes and cDCs were measured in additional falciparum malaria patients from the same study site at day 0 and day 28 (*n*=8) (**Fig. 2E, Fig. S2**). Consistent with transcriptional findings, the majority of individuals had reduced inflammatory cytokine/chemokine secretion in CD14^+^ monocytes at day 0 compared to day 28. This reduction was significant at the population level for MCP1 (**Fig. 2F**). MCP1 expression was also reduced at day 0 compared to day 28 in cDCs. To measure the co-expression of cytokines, responses were analysed by SPICE (Simplified presentation of incredibly complex evaluations(*46*)). Overall, the composition of cytokine expression in CD14^+^ monocytes at day 0 was significantly different to expression at day 28 (**Fig. 2G**). Collectively, data reveal a significant enrichment of regulatory innate cells during malaria infection which down-regulate multiple cytokine and chemokine responses along with HLA-DR associated genes and pathways required for robust inflammatory control and antigen presentation in a cell subset specific manner, regulated by cell subset specific pathways. In contrast, multiple receptors involved in antibody mediated functions are upregulated, consistent with a potential role of innate cells in antibody mediated parasite clearance during infection.

### Subset-specific activation and regulation of NK cells during malaria

Along with innate myeloid cells, NK cells are important early responders to *Plasmodium* infection, and have roles in adaptive immunity as effector cells. Broadly, NK cells exist in multiple distinct functional subsets along a spectrum of least differentiated CD56 bright cells, towards highly differentiated CD57^+^ senescent cells. CD56 bright NK cells and other less differentiated subsets produce IFNγ following parasite stimulation *in vitro* (*47, 48*) and during controlled human malaria infection (*49*). In contrast, adaptive and highly differentiated NK cells expand in malaria exposed individuals, and function via antibody dependent cellular cytotoxicity (ADCC) to protect from malaria (*50, 51*).

To investigate the transcriptional activation of these phenotypically distinct NK cell populations during malaria, we first undertook unbiased sub-clustering of NK cells identified in PBMCs (**Fig. 1**). Clustering identified five subsets which were annotated based on cluster markers as CD56 bright, Transitional, IFNγ^+^ Adaptive, IFNγ^-^ Adaptive and PD1^+^ NK cell subsets (**Fig. 3A, Table S4**). CD56 bright cells expressed the highest levels of *NCAM1* (encoding CD56), *SELL*, *KLRF1, GZMK* and *IL7R.* The Adaptive cell subset had increased expression of *NKG7, GZMB* and *FCGR3A* (encoding CD16). Transitional NK cells expressed markers from both CD56 bright and Adaptive subsets, consistent with previous scRNAseq analysis (*52, 53*)(**Fig. 3B**). Within the Adaptive cell subset, two clusters were further identified, differentiated as IFNγ^+^ and IFNγ^-^ Adaptive subsets based on *IFNG* expression, along with *TNF*, *CCL3* and *CCL4*. Additionally, we identified a NK cell cluster expressing high levels of *PDCD1* (encoding PD1). PD1^+^ NK cells have previously been shown to expand with age in malaria endemic populations and have increased function in ADCC (*10*). PD1^+^ NK cells also had increased expression of *VCAM1, ITGAD* (encoding CD11d)*, TOX, TNFRSF1B* (encoding TNFRII) and *CD160* (**Fig. 3B**). During malaria, there was a significant increase in the proportion of IFNγ^+^ Adaptive cells, consistent with an increased inflammatory and cytokine responsiveness of Adaptive NK cells during infection. Additionally, there was a proportional decrease in PD1^+^ NK cells during acute infection compared to 7 days following treatment (**Fig. 3C**). This decreased proportion of PD1^+^ NK cells during acute malaria is in contrast to previous reports of the expansion of this subset identified by flow-cytometry in Malians with falciparum malaria (*10*).

**Fig. 3.**
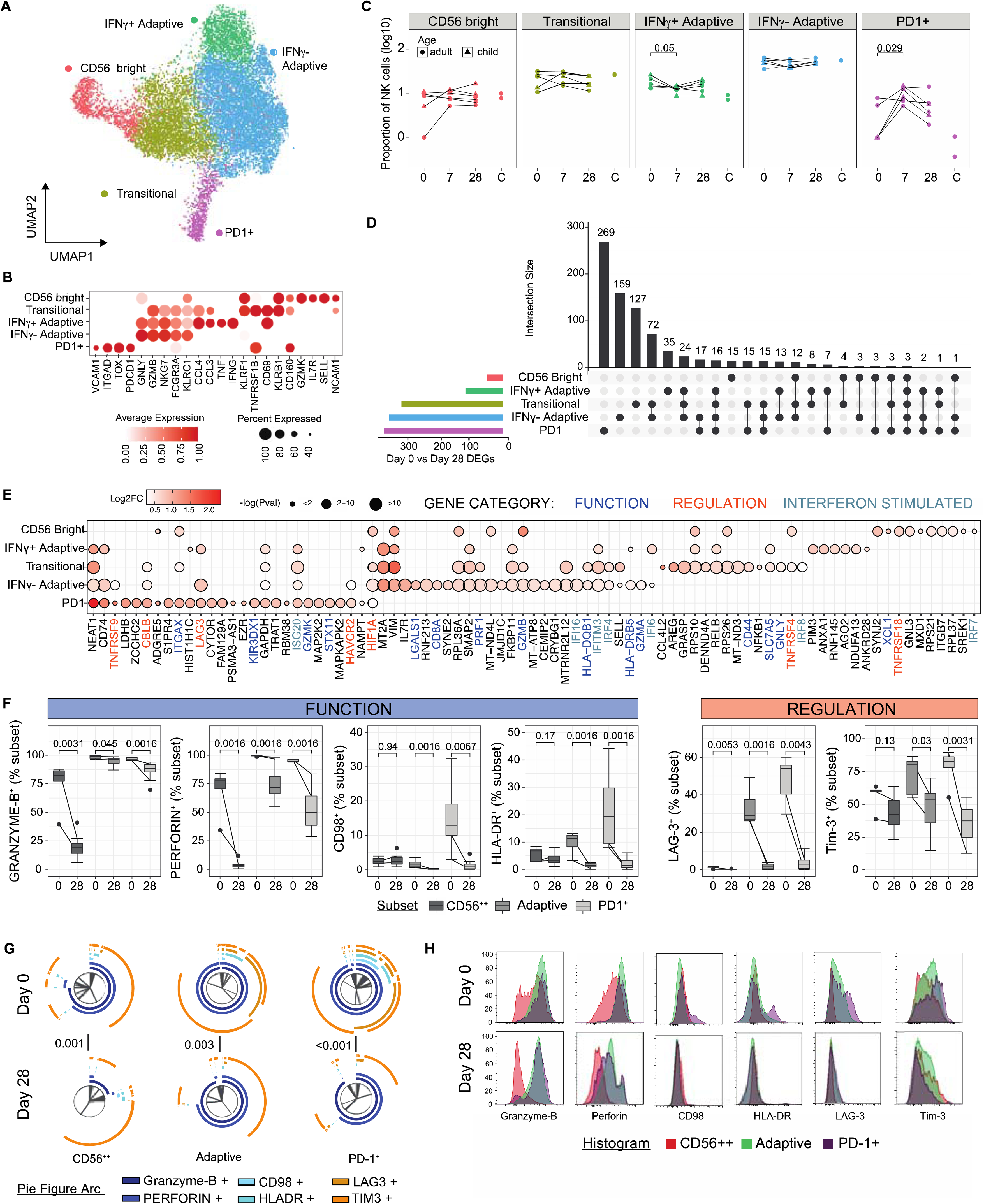
Subset specific activation and regulation of NK cells in malaria. (**A**/**B**) Five subsets of NK cells were identified based on unsupervised clustering and marker expression as CD56 bright, Transitional, IFNγ^+^ Adaptive, IFNγ^-^ Adaptive and PD1^+^ subsets. (**C**) Relative proportion of identified subsets during malaria infection (day 0), 7- and 28-days post treatment, and in healthy uninfected individuals. P-value indicated is calculated by Mann-Whitney U test between day 0 and indicated subsequent time points (**D**) Upset plot of DEGs in NK cell subsets day 0 compared to day 28. The number of shared and subset specific DEGs indicated. (**E**) Top 20 DEGs of in each NK subsets, and additional genes of interest. Genes with known roles in regulation and/or function are indicated. (**F**) PBMCs from individuals with P. falciparum malaria (Day 0, *n* = 5, patients 28-days post-infection (Day 28, *n* = 8) were analysed ex vivo to detect NK cell protein expression of identified genes by flow cytometry. Expression of proteins related to function and regulation of NK cells, shown as positive frequencies of NK cell subsets CD56 bright (CD56^++^), Adaptive and PD1^+^. Box plots show the median and IQR of volunteers, lines represent paired observations, group comparisons performed by Mann-Whitney U test. (**G**) Co-expression of proteins related to function and regulation of NK cells analysed by SPICE. Pie graphs comparisons performed by Permutation test. (**H**) Histograms normalised to mode show expression of gene-related proteins within NK cell subsets of concatenated group data. See also Fig. S3-S4 and Table S4-S5.

To investigate malaria-driven transcriptional changes to NK cells, we identified DEGs comparing day 0 to day 28 within each subset. Transitional, IFNγ^-^ Adaptive and PD1^+^ subsets were more transcriptionally active during malaria compared to CD56 bright and IFNγ^+^ Adaptive subsets (**Fig. 3D, Table S5**). PD1^+^, Transitional and IFNγ^-^ Adaptive NK cells, had a large proportion of subset specific DEGs, suggesting unique NK cell subset specific activation pathways during malaria (subset specific DEGs 76%, 42% and 47% respectively, **Fig. 3D**). During acute malaria, there was evidence for increased cytotoxic potential across multiple NK cell subsets, with upregulation of genes with known cytotoxic functions (Granzyme members *GZMB, GZMA, GZMK* (*54*), Granulysin *[GNLY],* Perforin *[PRF1])* and roles in degranulation and NK cell activation (*CD44* (*55*)*, CCL4L2* (*56*)*, STX11* [encoding Syntaxin 11 (*57*)], *CD8A*(*58*)*, LGALS1* [encoding Galectin 1 (*59*)], *XCL1, SLC7A5* [encoding CD98/LAT1 (*60*)] and *SELL* [encoding CD62L (*61*)]) (**Fig. 3E**). GSEA confirmed upregulation of multiple pathways associated with cell function, particularly within Transitional, IFNγ^-^ Adaptive and IFNγ^+^ Adaptive subsets (**Fig. S3**). In PD1^+^ NK cells, there was also evidence for subset specific upregulation of the MAPK pathway (increased *MAPKAPK2* and *MAP2K2*). Additionally, PD1^+^ NK cells had a large increase in *ITGAX* (encoding CD11c) expression during acute infection, with smaller increases in *ITGAX* in CD56 Bright and Transitional NK cells. CD11c expression in NK cells is upregulated in response to inflammatory cytokines (*62*), and is a marker of bitypic NK cells which can produce inflammatory cytokines and also drive the proliferation of γδ T cells (*63*). Further, upregulation of HLA-DR genes *HLA-DQB1* and *HLA-DRB5* were also detected, with HLA-DR previously associated with NK cell activation and antigen-presentation in some settings (*64*).

Upregulation of genes mediating increased cytotoxicity and function during malaria was balanced by increases in multiple negative regulators of NK cells, including *TNFRSF4* (encoding CD134/OX40(*65*)), *TNFRSF9* (encoding CD137/4-1BB (*66*)), *TNFRSF18* (encoding CD357/GITR (*67*)), *LAG3* (*68*)*, HAVCR2* (encoding Tim-3 (*69*)) and *CBLB* (*70*). GSEA showed significant enrichment of negative regulatory pathways in PD1^+^ and IFNγ^-^ Adaptive subsets (‘negative regulation of immune system process’ and ‘negative regulation of lymphocyte activation’)(**Fig. S3**). Across all subsets, multiple type I IFN signalling genes were upregulated including *ISG20, IFI16, IFI6, IFITM3, IRF4, IRF8,* and *IRF7* and GSEA confirmed upregulation of ‘response to type I IFN’ in Transitional and IFNγ^-^ Adaptive cells (**Fig. S3**). Type I IFN signalling in NK cells has been shown to suppress IFNγ production during viral infection (*71*). Taken together data suggest that Type I IFN signalling is activated in NK cells in response to malaria, which may act as a regulatory of NK cell inflammatory response during infection. To investigate these key transcriptional changes to NK cells during malaria at the protein level, we analysed NK cells by flow cytometry in additional falciparum malaria patients (day 0) and patients 28 days after infection. NK cells were identified as CD56 bright, Adaptive and PD1^+^ subsets (*10*)(**Fig. S4A**). Within these patients, there was a slight increase in CD56 bright cells within the NK compartment during infection (**Fig. S4B**). Across all three NK subsets, there was a significant increase in Granzyme B and Perforin expression at day 0 compared to day 28 post treatment (**Fig. 3F**). In contrast, CD98 and HLA-DR expression were increased on Adaptive and PD1^+^ NK cells, but not CD56 bright NK cells, and expression levels were much higher on PD1^+^ cell subset. Similarly, the regulatory marker LAG-3 was increased in expression on all NK cell subsets at day 0 (at very low levels on CD56 bright cells), but Tim-3 was only increased on Adaptive and PD1^+^ cells (**Fig. 3F**). IFNγ and TNF expression were also assessed. While *ex vivo* IFNγ expression was below the limit of detection for all NK cell subsets, there was a significant increase in TNF production in Adaptive, but not CD56 bright NK cells during malaria (**Fig. S4C**). Additionally, there was a significant increase in CD11c expression on Adaptive NK, but not CD56 bright cells (**Fig. S4D**). When considering the total profile of Granzyme B, Perforin, CD98, HLA-DR, LAG-3 and Tim-3 on the three distinct NK cell subsets, there was a significant difference in the overall composition of marker expression in each subset between day 0 and day 28, indicating significant upregulation of both cytotoxic and regulatory markers during infection (**Fig. 3G**). However, PD1^+^ NK cells are the most highly activated and regulatory during malaria due to the increased overall level of expression of these markers during acute infection, particularly CD98, HLA-DR, LAG-3 and Tim-3, compared to adaptive and CD56 bright NK cells (**Fig. 3H, Fig. S4E**)

### Activation and regulation of γδ T cells with diverse functions during malaria

γδ T cells are key innate cell responders during malaria which proliferate in response to malaria parasites and produce inflammatory cytokines with important roles in protection (*72*). γδ T cells can also recognize and kill parasites via lysis, and opsonic phagocytosis(*73*), and present antigen to activate T cells (*74*). However, in individuals who have had repeated malaria infections, γδ T cells become tolerized, with reduced cell frequency, inflammatory responses and increased expression of regulatory proteins (*11, 75*). γδ T cells in highly exposed individuals express increased CD16 and have increased cytolytic responsiveness to opsonized parasites (*76*). To explore these multiple roles and tolerization mechanisms of γδ T cells in the patients in this study, we sub-clustered γδ T cells identified in PBMCs (**Fig. 1**) and categorized 6 clusters as Cytotoxic, Inflammatory, Antigen-presenting, Transitional, Type 3 and Naive γδ T cells (**Fig. 4A, Table S6**). Cytotoxic subset cells expressed the highest levels of *GNLY*, *GZMB*, *GZMH*, *NKG7*, *FCGR3A* (encoding CD16) and *FGFBP2;* Inflammatory subset cells expressed high levels of *CCL4L2*, *CCL4*, *CCL3*, *IFNG* and *TNF*; Antigen-presenting subset cells expressed HLA-II related genes; and Transitional γδ T cells were characterized by expression of genes that drive the early differentiation of T cells *IL21R*, *IER5L*, *YPEL5* and *IFRD1* (**Fig. 4B**). We also identified Type 3-like γδ cells with high expression of *KLRB1*, *NCR3*, *RORA* and *IL7R*, and Naive γδ T cells with high expression of *LTB*, *CCR7* and *AQP4*, as in previous scRNAseq data sets (*77*). Within γδ T cells, there was an expansion of the Cytotoxic cell subset at day 7 and 28 after infection, and expansion of the Transitional cell subset at day 7 (**Fig. S5A**).

**Fig. 4.**
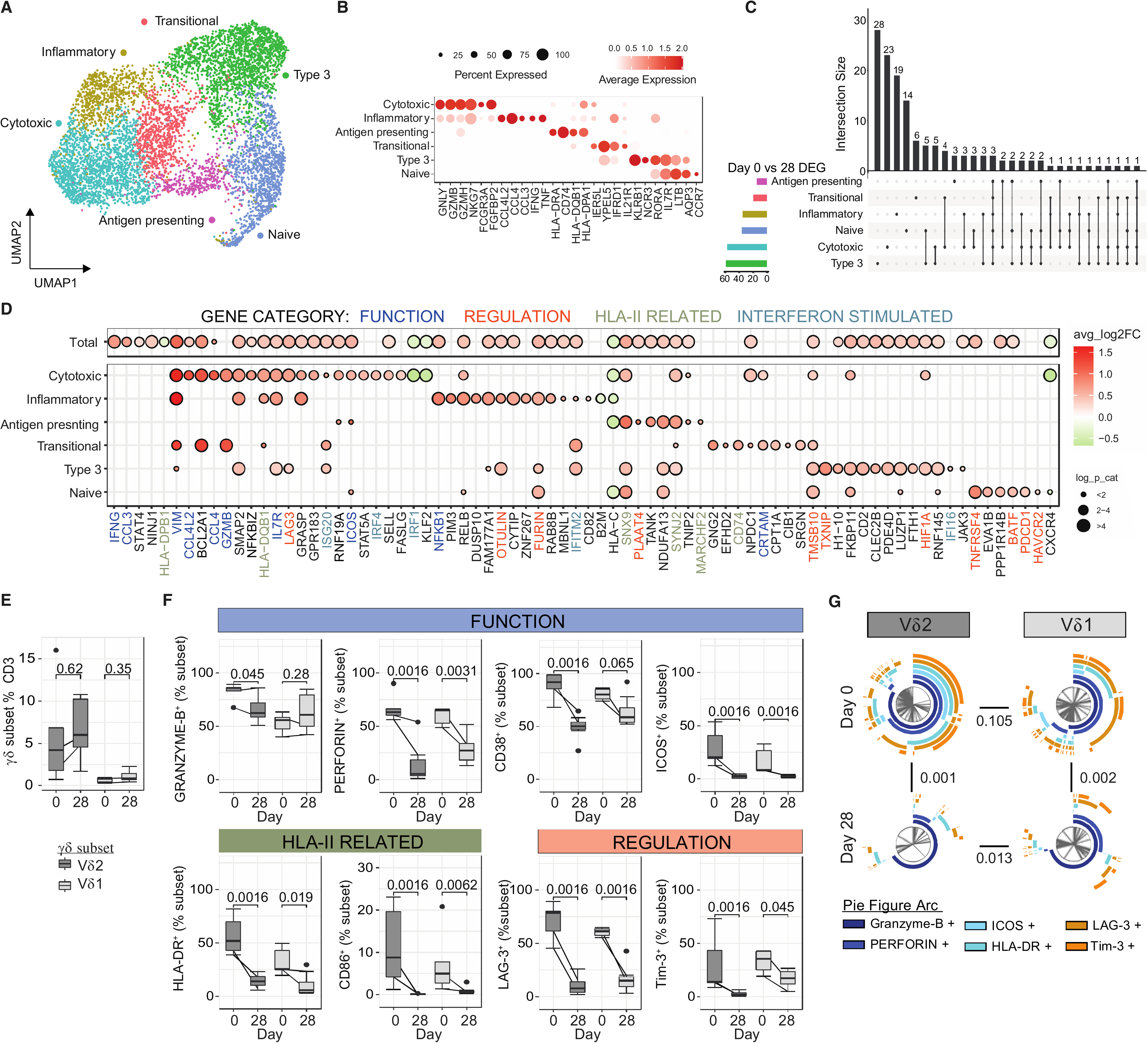
Inflammatory activation and regulation of effector γδ T cells during malaria. (**A**/**B**) Five subsets of γδ T cells were identified based on unsupervised clustering and marker expression as Cytotoxic, Inflammatory, Antigen-presenting, Transitional, Type 3 and Naive. (**C**) Numbers of upregulated DEGs in γδ T cell subsets day 0 compared to day 28. The number of shared and subset specific DEGs indicated. (**D**) Top 20 upregulated DEGs in each γδ T cells subsets, and additional genes of interest. Genes with known roles in regulation and/or function are indicated. (**E**) PBMCs from patients with falciparum malaria (Day 0, *n* = 5) and convalescent malaria patients 28-days post-infection (Day 28, *n* = 8) were analysed ex vivo to detect Vδ2 and Vδ1 γδ T cells and measure protein expression of identified genes by flow cytometry. (**F**) Expression of proteins related to function and regulation of γδ T cells, shown as positive frequencies of Vδ2 and Vδ1 γδ T cells. Box plots show the median and IQR of volunteers, lines represent paired observations, P values indicated are calculated by Mann-Whitney U test. (**G**) Co-expression of proteins related to function and regulation of Vδ2 and Vδ1 γδ T cells analysed in SPICE. Pie graphs comparisons performed by Permutation test. See also Fig. S5 and Tables S6-7.

To investigate γδ T cell transcriptional changes during malaria, DEGs were identified in γδ T cell subsets, and due to low cell numbers in subsets, also the total γδ T cell population (**Fig. 4C-D, Table S7**). Both shared and subset specific transcriptional changes between day 0 and day 28 where identified (**Fig. 4C-D**). DEGs included upregulation of inflammatory cytokines/chemokines *IFNG, CCL3, CCL4, CCL4L2* and genes associated with T cell activation (including *VIM*, *ICOS*, *IL7R* (encoding CD127)) across multiple subsets, consistent with the well documented inflammatory responsiveness of γδ T cell during malaria infection (*72*). Additionally, consistent with increased cytotoxic capacity during malaria reported previously (*73*), expression of cytotoxic serine protease *GZMB* was increased in both Cytotoxic and Transitional γδ T cells during malaria. Further, *CRTAM,* which drives the development of cytotoxic CD4 and CD8 T cells (*78*), was increased during infection, consistent with the expansion of Cytotoxic γδ T cell subsets at days 7 and 28 (**Fig. 4D)**. Along with increased inflammatory and cytotoxic capacity, there was a suggestion of increased antigen presenting properties during infection, with upregulation of HLA-II related genes (*CD74, HLA-DPB1, HLA-DQB1*), and genes related to endocytosis and intracellular vesicle trafficking (such as *SNX9* and *SYNJ2*). Together, these data are is indicative of polyfunctional γδ T cell activation during infection, consistent with previous phenotypic data (*72–74*).

Along with activation of multiple γδ T cell functions, increased expression of genes with roles in regulation and cell exhaustion were detected (**Fig. 4D**). These included upregulation of *LAG3* on Cytotoxic γδ T cells and increased expression of inflammatory regulators *OTULIN* (*79*) and *FURIN* (*80*) on Inflammatory γδ T cells. Within Type 3 and Naive γδ T cells, upregulated genes included those related to inflammatory regulation (*TMSB10;* which suppresses inflammatory macrophages)(*81*), *HIF1A;* which controls γδ T cell mediated inflammation(*82*) and *BATF;* which is upregulated in exhausted CD8 T cells (*83*)), cell exhaustion (*PDCD1* (encoding PD1); which dampens inflammatory and cytotoxic potential of γδ T cell (*84, 85*), *TNFRSF4* (encoding OX40) and *HAVCR2* (encoding Tim-3); which reduces cytokine and cytotoxic potential of γδ T cells (*86*) and pro-apoptotic signalling (*TXNIP*). Similar to myeloid and NK cell responses, DEGs included upregulation of multiple Type I IFN response genes including *ISG20, IRF4, IRF1, IFITM2, IFI16*. γδ T cells have been reported to respond to Type I IFNs produced from poly(I:C) activated cDCs in other settings (*87*). However, to the best of our knowledge, a role of Type I IFNs in driving tolerogenic γδ T cells has not been explored.

To confirm our transcriptional findings, we assessed protein-level expression of multiple functional and regulatory markers in additional patient samples, by FACS (**Fig. S5B**). While we were unable to differentiate between Vδ2 and Vδ1 γδ T cells in our 3’ transcriptional data set, the large majority of circulating γδ T cell within the CD3 T cell compartment at both day 0 during acute infection and day 28 post treatment were Vδ2 γδ T cells (**Fig. 4E**). Within Vδ2 cells γδ T cells, there was increased expression of Granzyme-B at day 0, but not in Vδ1 γδ T cells as detected via FACS (**Fig. 4F)**. Other functional and activation markers, Perforin, CD38, ICOS, and HLA-DR were increased on both Vδ2 and Vδ1 γδ T cells during malaria. Similarly, both LAG-3 and Tim-3 were increased in expression on both subsets (**Fig. 4F**). Granzyme-B, ICOS and HLA-DR expression was higher on Vδ2 compared to Vδ1 γδ T cells during malaria (**Fig. S5C**). To understand the expression of different functional, activation and regulatory markers on Vδ2 and Vδ1 γδ T cell subsets, marker expression was analysed by SPICE. Both Vδ2 and Vδ1 γδ T cells had high levels of co-expression of key proteins related to multiple functions and regulation (**Fig. 4G**). The magnitude and composition of marker co-expression by γδ T cells was increased at day 0 compared to day 28 expression. Together data shows that, as seen in myeloid and NK cells, Type I IFN signaling is activated in γδ T cells during malaria, and γδ T increase both functional and regulatory functions during malaria.

### CD4 T cell response is dominated by Type 1 regulatory cells during malaria which share signatures with Th1 cells

CD4 T cells play multiple essential roles in protection from malaria, including IFNγ mediated direct-killing of parasites, and by providing help to B cells to produce antibodies required for protection (*88*). However, multiple lines of evidence have shown that malaria drives the expansion of regulatory CD4 T cells, particularly Type 1 regulatory (Tr1) cells that co-produce IFNγ and IL10 in this disease. These cells rapidly expand via Type I IFN signalling in initial parasite infection in humans (*16*), and dominate the parasite specific CD4 T cell response in children in endemic areas (*13–15*). To investigate CD4 T cells transcriptionally during malaria, we subclustered CD4 T cells from PBMCs, with 11 subsets identified (**Fig. 5A**). These CD4 T cell subsets were annotated as naive, activated, T-regulatory (Treg), T-follicular helper (Tfh), T-follicular regulatory (Tf-reg), T-helper (Th) 1, Tr1, Th2 and Th17 subsets based on expression of T naive, activation, Treg and T helper signatures (*89–92*) and canonical marker genes (**Fig. 5B, Table S8**). The proportions of each subset within the CD4 T cells compartment did not significantly change between acute infection and post treatment timepoints. (**Fig. S6)**. We conducted DEG analysis for each CD4 T cell subset, on day 0 (malaria) compared to day 28 (post treatment). Tr1 cells and Th1 CD4 T cells were highly transcriptionally active during infection, while Tregs and Tfregs were the least activated (**Fig. 5C, Table S9)**. In Tr1 cells, 68% of DEGs were unique, consistent with a Tr1 specific activation program during malaria (**Fig. 5C**). Upregulated genes included canonical markers of *IFNG* and *IL10*, and many other genes with known roles in immunosuppression and/or function of regulatory T cells including *LAG3*, *HAVCR2* (encoding Tim-3 (*93*)), *TNFR2* (*94*), *CTLA4* (*95*), *TNFRSF4* (encoding OX40/CD134 (*96*)), *TNFRSF18* (encoding GITR/CD375 (*97*)), *PDCD1* (encoding PD1) and *CCL4* (*98*) (**Fig. 5D**). The upregulation of multiple co-inhibitory receptors is consistent with our recent data showing Tr1 cells express overlapping co-regulatory proteins(*19*). Additionally, *IKZF3* which has been shown to have high expression in IL10^+^ CD4 T cells(*99*), and *LAIR2* which has been identified as a core Treg signature gene in humans(*100*) had increased expression on Tr1 cells during malaria (**Fig. 5D**). Tr1 cells also showed significant activation during infection, with marked upregulation of *CD38* and *ICOS* during infection. *CD38* upregulation was unique to Tr1 cells, while *ICOS* was also increased on Th1 cells.

**Fig. 5.**
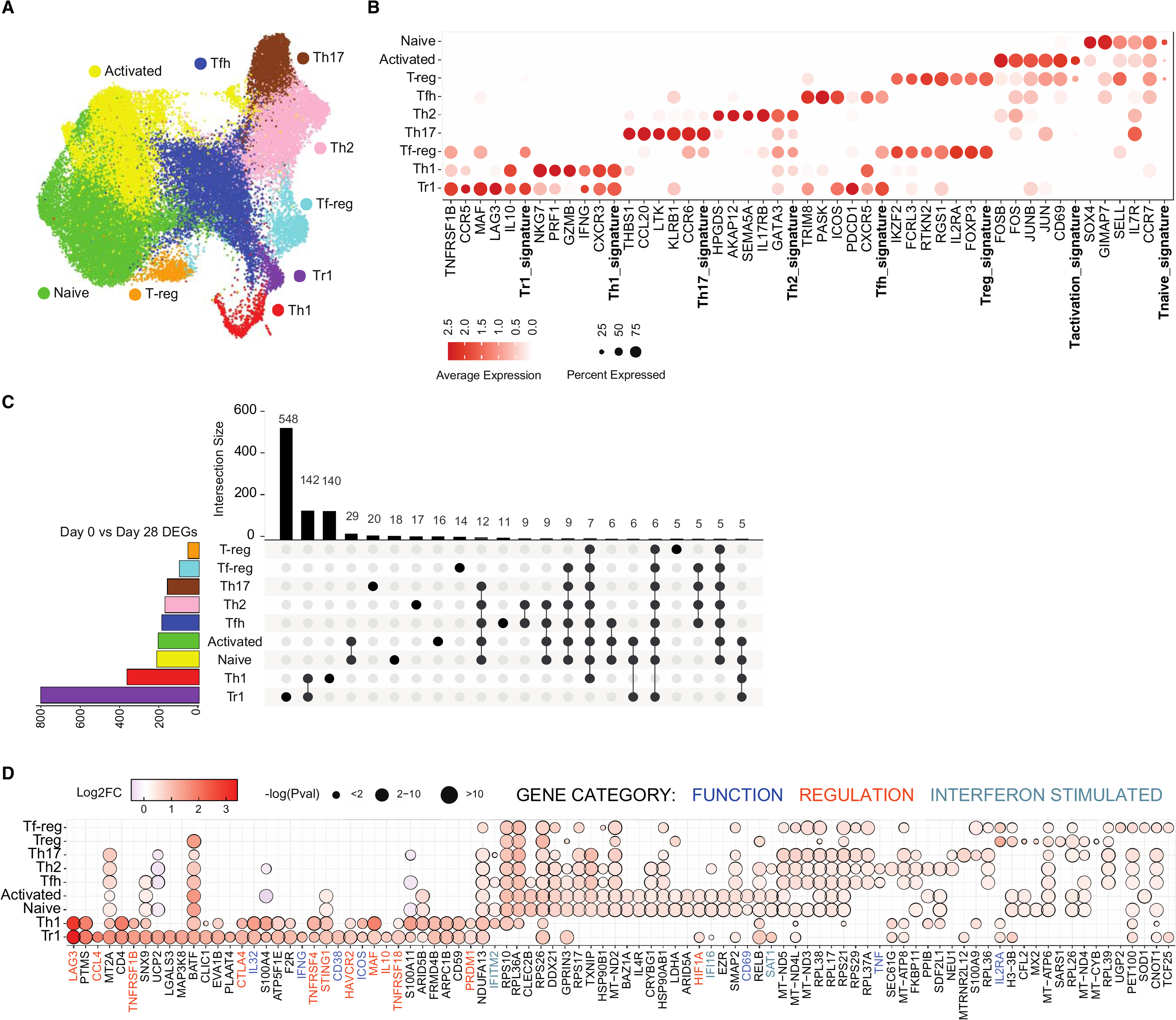
Tr1 CD4 T cells dominate the response during malaria. (**A**/**B**) Subsets of CD4 T cells were identified based on unsupervised clustering and categorized based on canonical marker expression and T helper expression signatures. (**C**) DEGs in CD4 T cell subsets at day 0 compared to day 28 were identified. The number of shared and subset specific DEGs indicated in Upset plot. Overlaps of <5 genes not shown. (**D**) Top upregulated 20 DEGs in CD4 T cell subsets. Genes with known roles in regulation and/or function are indicated. See also Fig. S6 and Tables S8-9.

Tr1 cells can emerge from Th1 cells that gain IL10 expression (*101*). Consistent with this, most of the top upregulated genes in Tr1 cells were shared with Th1 cells, including regulatory markers *LAG3, TNFRSF4* (encoding OX40), *TNFRSF1B* (encoding TNFR2), *TNFRSF18* (encoding GITR), *HAVCR2* (encoding Tim-3) and *CTLA4* (**Fig. 5D**). Additional upregulated genes in Th1 cells included *MAF,* which drives IL10 expression in Th1 cells in murine malaria models (*102*), *PRDM1* (encoding BLIMP1), which promotes IL10 in Tr1 cells in murine malaria models (*103*) and is highly expressed in malaria-specific Tr1 cells in Ugandan children (*13*), and *STING1* (encoding STING – stimulator of interferon response cGAMP interactor 1), which we have recently shown to be a central driver of Tr1 cell development (*104*). In Th1 and/or Tr1 cells, several other Type I IFN responses genes were up regulated, including *IFITM2, IFI16, SAT1, IFI35, IFI27L2, LYE6* and *ISG15* (**Fig. 5D, Table S9**). In other CD4 T cell subsets, the number of DEGs and magnitudes of fold changes to expression were relatively lower, with many ribosomal and mitochondrial genes shared across non-Th1/Tr1 subsets (**Fig. 5C-D**). However, *BATF,* which is critical for Th17 and Tfh cell differentiation(*105, 106*) was upregulated across all CD4 T cell subsets except for FoxP3^+^ Tregs. While the roles of Th17 cells in human malaria are largely unknown, Tfh cell activation and development is critical for the induction of humoral responses required to drive protection against malaria (*107*). Tfh cell activation during malaria is skewed towards Th1-Tfh cell responses (*108–110*), and consistent with this, Th2 and Tfh cells subsets upregulated both the Th1 associated cytokine TNF, and *ETS1* which represses Th2-Tfh subset differentiation in both human and mouse models of systemic lupus erythematosus (*111*). *ETS1* is also essential for *BATF* function in effector T cells (*112*). While DEGs of Tregs and Tf-regs were the lowest of all subsets, *IL2RA*, essential for Treg function(*113*), was upregulated in malaria in both subsets, indicative of increased functional Tregs and Tf-reg cells during infection. Together these data are indicative of CD4 T cell activation dominated by Tr1 cells with increases suppressive function during malaria, and the emergence of Tr1 cells from Th1 cells via Type I IFN signalling(*16, 19*).

### Expansion of IL10^+^ regulatory B cells during malaria infection

Malaria specific B cell responses are essential for development of immunity against malaria, with antibodies being key mediators of protection through control of parasite burden(*114*). While robust memory B cells and sustained antibodies can develop against malaria(*22*), there is evidence that these responses may also be negatively impacted by malaria driven immunomodulation. Memory B cell responses are suboptimal in some malaria transmission settings, with lower levels of antibody production, short-lived antibody responses and expanded ‘atypical’ memory B cells (*20–22*). Atypical memory B cells express high levels of FCLR5, and appear to have reduced functional capacity compared to ‘typical’ memory B cells (*23, 24*). However, ‘atypical’ responses emerge in both infection and vaccination (*115*), mount recall responses (*116*) and can produce antibodies with the support of T-follicular helper cells (*117*). Therefore, whether atypical memory B cells are protective or disruptive in protective immunity remains unclear.

To investigate transcriptional changes to B cells during malaria within our PBMC data set, B cells were sub-clustered to identify 7 subsets of B cells and annotated as transitional, naive, memory, activated, atypical, plasmablast and proliferating plasmablast populations (**Fig. 6A**). Based on previously published studies (*115, 118, 119*), identified subsets included transitional B cells with high expression of *MME (*encoding CD10) and naive B cells with high expression of *TCL1A*, *NEIL1, IGHD*, *IGHM*, *FOXP1, BACH2* and *IL4R;* quiescent memory B cells which had relatively high expression of memory marker *CCR7*, and upregulated *MARCKS, CD82, IER5, CD70 and CD80* which are increased in expression on memory relative to naive B cells in previously published data sets(*120, 121*) annotated by the Human Protein Atlas (proteinatlass.org); activated B cells with increased expression of *CD1C, CD79B, ACTG1, GAPDH, TKT, S100A10, CRIP2, CPNE5,* and *GSN;* ‘Atypical’ memory B cells with high expression of *FCLR5, FCLR3, ITGAX* (encoding CD11c)*, TNFRSF1B* and *CD86;* plasmablasts with expression of *MZB1, XBP1, IRF4, PRDM1 (*encoding BLIMP1), and switched IgG genes; and proliferating plasma blasts which expressed plasmablast marker genes along with high levels of *MKI67* (**Fig. 6B**, **Table S10**). During infection (day 0), there was an increased proportion of plasmablasts, which made up to 20% of the B cell compartment, consistent with previous studies of in Ugandan (*122*) and Kenyan children with malaria(*123*), and in adults during controlled human malaria infection (*124*) (**Fig. S7A**).

**Fig. 6.**
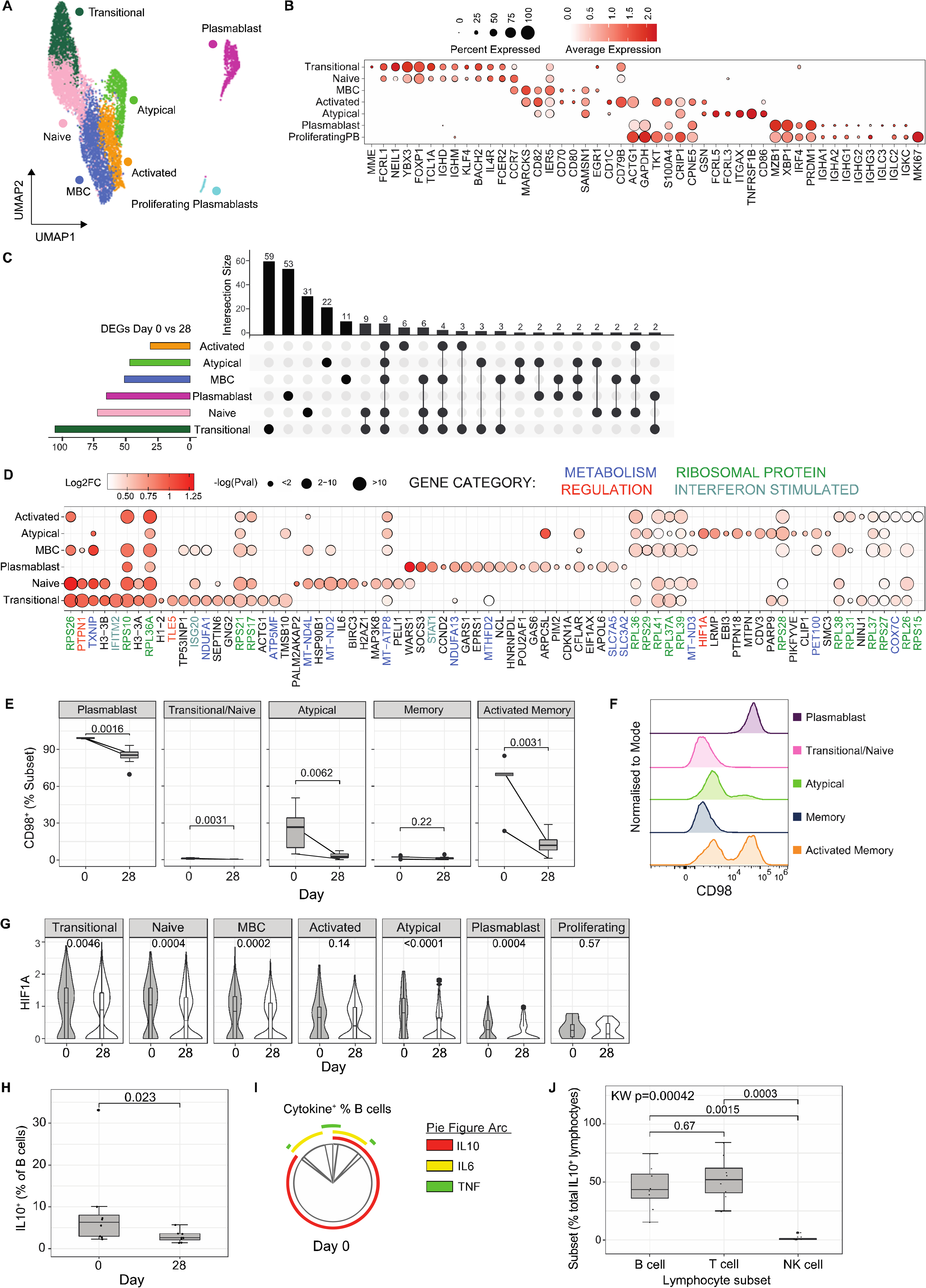
B cell activation and induction of IL10^+^ Bregs during infection. (**A**/**B**) Subsets of B cells were identified based on unsupervised clustering and categorised based on marker expression. (**C**) DEGs in B cell subsets day 0 compared to day 28 were identified. The number of shared and subset specific DEGs indicated in Upset plot. (**D**) Top upregulated 20 DEGs in B cell subsets. Genes with known roles in regulation and/or function are indicated. (**E**) CD98 protein expression was quantified on plasmablasts, transitional/naive, atypical, memory and activated memory B cell subsets at day 0 (*n* = 5) and day 28 (*n* = 8). Box plots show the median and IQR of volunteers, lines represent paired observations, group comparisons performed by Mann-Whitney U test. (**F**) CD98 MFI of concatenated samples at day 0 (*n* = 5). (**G**) *HIF1A* mRNA expression in each B cell subset at day 0 and 28. P-value calculated by Mann-Whitney U test P is indicated (unadjusted). (**H**) IL10 protein expression on B cells during malaria (day 0, *n* = 8) and day 28 post treatment (*n* = 8), Wilcoxon rank sum test is indicated. (**I**) Co-expression of B cell IL10 production with IL6 and TNF during malaria (*n* = 8) analysed in SPICE. (**J**) The proportion of each lymphocyte subset contributing to IL10 lymphocyte production during malaria (day 0, *n* = 8). P-value calculated using Kruskal wallis and post-Dunn test (FDR adjusted) indicated.

DEG analysis of each B cell subset comparing day 0 to day 28, identified large numbers of DEGs across all subsets except proliferating plasmablasts (**Fig. 6C**, **Table S11**). DEGs were both shared and subset specific, and a large number of the top DEGs for each subset were ribosomal proteins, possibly indicating increased protein synthesis and highly activated states of B cells during infection (**Fig. 6D**). Additionally, many DEGs had roles in metabolism, consistent with reshaping of energy use and metabolic programs during B cell activation (*125*) (metabolic associated DEGs include members of NADH dehydrogenase complex *NDUFA1, NDUFA13, MT-ND4L, MT-ND2, MT-ND3*, cytochrome C oxidase subunit *COX7C* and chaperone *PET100*, ATP synthase subunits *ATP5MF* and *MT-ATP8).* Across B cell subsets, multiple upregulated Type I IFN signaling response genes were detected, including *IFNITM2, ISG20,* and *STAT1*, consistent with the important role of Type I IFN signaling in malaria immune responses across multiple cell subsets. There was evidence that metabolic remodeling during infection was B cell subset specific. For example, transitional, naive, memory B cells and atypical memory B cells during acute infection had increased expression of *TXNIP,* a glucose feedback sensor which inhibits glucose uptake (*126, 127*). In contrast, plasmablasts had increased *SLC7A5* (encoding LAT1) and *SLC3A2* (encoding CD98 which interacts with LAT1 to transport L-glutamine), consistent with our recent findings of the importance of plasmablasts as negative regulators of germinal centre development via acting as a nutrient sink in mice models of malaria (*124*) (*128*). To confirm upregulation of glutamine transport on plasmablasts, CD98 levels were measured across B cell subsets in additional patients. We assessed CD98 expression during and after malaria in plasmablasts (CD27^+^CD38^+^), transitional/naive IgD^+^ B cells, and IgD-B cell subsets (atypical [CD27-CD21-], memory [CD27^+^CD21^+^] and activated memory [CD27^+^CD21-]) (**Figure S7B-C**). In these additional participants, plasmablasts made up to 30% of the B cell compartment during malaria, but less than 2% at 28 days post-treatment (**Fig. S7D**). There was also a significant increase in the proportion of activated memory cells during malaria (**Fig. S7D**). The frequency of CD98^+^ cells increased during malaria in plasmablasts, transitional/naive, atypical and activated memory B cells (**Fig. 6E**). However, the frequency of CD98^+^ cells, and the magnitude of CD98 expression was far greater on plasmablasts compared to other subsets (**Fig. 6F**, **Fig S7E-F**). Together, these data are consistent with a potential negative role of plasmablast expansion and CD98 expression as a nutrient sink that limits productive germinal center activation during infection(*124*), but shows that activated memory B cells also upregulate glutamine transport during infection.

Along with a potential disruptive role of plasmablasts in malaria infection, several upregulated DEGs where suggestive of other tolerized/immunosuppressed B cell responses during infection. For example, *PTPNI* (encoding PTP1B) which was upregulated in transitional, naive and memory B cell subsets, negatively regulates B cell signally via CD40 and BAFF-R and TLR4, and downregulates T-dependent immune responses (*128*). *TLE5* (also known as AES), which negatively regulates NF-κβ signalling (*129*), required for B cell activation and survival (*130*), was upregulated on transitional B cells during infection. Of note, *HIF1A,* which drives B cell IL10 production in hypoxic conditions (*131*), was significantly upregulated in Atypical B cell during acute infection. Further interrogation of *HIF1A* suggested increased expression during acute infection also occurred in transitional, naive, memory B cells and plasmablasts subsets (**Fig. 6G**), consistent with the capacity of diversity of human B cells subsets to produce IL10 (*132*). IL10 production by B cells is indicative of Breg subsets, which have been shown in mice to be a major source of IL10 during infection, and protect from experimental cerebral malaria (*25, 133*), however have yet to be identified during human malaria. To confirm the expansion of IL10^+^ Bregs during falciparum malaria, we quantified IL10^+^ production in B cells in additional study participants. We analyzed the total B cell population, due to the diverse B cell phenotypes of IL10^+^ Bregs, and measured TNF and IL6 production, which are often co-produced with IL10(*132*) (**Fig. S8A**). We confirmed that there was significant increase in IL10 production in B cells during malaria, indicating of malaria induction of Bregs (**Fig. 6H**). In contrast, there was no evidence for increased B cells expression of IL6 nor TNF, despite increased IL6 transcripts levels (**Fig. S8B**). Although a previous study reported co-expression of TNF and IL6 by IL10 producing Bregs (*132*), only a small fraction of IL10^+^ Bregs co-produced IL6 during malaria, and there was minimal co-expression with TNF (**Fig. 6I**). To assess the relative importance of Bregs, compared to IL10^+^ T cells (largely Tr1 CD4 T cells), the proportion of B cells amongst all IL10 producing lymphocytes was measured. The proportion of IL10 lymphocytes that were Bregs was comparable to CD3 cells as the source of IL10 from lymphocytes (**Fig. 6J**), identifying IL10 Bregs as a potentially important contributor to the regulatory/tolerogenic response during malaria infection in humans.

## DISCUSSION

Malaria drives tolerogenic immune cell responses which protect from inflammation mediated immunopathogenesis at the costs of reduced parasite control and suboptimal adaptive immunity. Here, using scRNAseq analysis of PBMCs during and following falciparum malaria, we comprehensively map malaria associated tolerogenic responses across innate and adaptive immune cells. These data show that malaria driven immunomodulation occurs across the immune landscape, with subset specific activation and regulatory programs identified. By analysing malaria-driven transcriptional changes at the subset level, we not only increase our understanding of how malaria modulates specific immune cell subsets, but also identify IL10^+^ Bregs as a major tolerogenic response in human adaptive immune cells during infection.

The use of scRNAseq analysis of PBMC immune responses during other infections has identified key protective and disrupted responses in various disease states (*28–31*). Here, we leveraged a large data set of >100 000 cells to understand malaria driven immune responses within major immune subsets, but also within subclustered cells. This approach allows for a granularity of understanding of specific cell responses not previously possible with bulk-level analysis. Within innate myeloid cells, malaria drove changes to monocytes consistent with the induction of immunosuppressive MS1-like monocytes, which have high expression of alarmins *S100A8/A9*, along with *RETN* and *ALOX5AP* and reduced expression of MHC class II. These immunosuppressive monocytes have been identified in scRNAseq data sets in both sepsis and COVID-19 patients (particularly those with severe disease), but not HIV infected individuals (*29, 30, 38*). During sepsis and COVID-19, immunosuppressive monocytes appear to emerge directly from inflammation-induced myelopoiesis within the bone marrow (*38, 134*). This pathway may also be important in malaria, with parasite infection shown to drive emergency myelopoiesis in mouse models (*135*). How these immunosuppressive monocytes protect from parasite-mediated immunopathology is unknown, however, the importance of tolerized monocytes in anti-disease immunity to malaria has been suggested by others (*8*). This anti-disease protection may come at a cost to both adaptive immunity by disruption of antigen presentation via down regulation of HLA-DR, which was also seen in DCs (*5*), and more broadly. For example, in sepsis, immunosuppressive monocytes have reduced responsiveness to LPS (TLR4 stimulation), consistent with dysregulated response to future bacterial infection in these patients (*38*). Reduced responsiveness to LPS has also been reported for monocytes exposed to *P. falciparum* parasites *in vitro* (*8*). As such, immunosuppressive monocytes may also be an important factor in the increased risk of bacterial infection in children with recent or acute malaria (*136*).

Evidence of malaria induced immunosuppression within our data was also observed in NK and γδ T cells, where multiple co-inhibitory receptors were upregulated during infection (including *TNFRSF4, TNFRSF9, TNFRSH18, HAVCR2, LAG3,* and *PD1*). Co-inhibitory receptors play important roles in regulating immune response to chronic infections, including malaria (*137*). For NK cell responses, increased PD1 expression has been previously reported in malaria exposed individuals previously (*10*), however the roles of other co-inhibitory receptors in regulating NK cell responses during malaria is less characterised. We have recently shown that a CD56neg NK cell subset is expanded in areas of high malaria burden and have important functional roles in protecting from diseases via antibody dependent cellular cytotoxicity (*138*). These CD56neg NK cells had high expression of LAG-3, a molecule which has been shown in other studies to be expressed on NK cells with increased glycolytic activity and to negatively regulate NK cytokine production but not cytotoxic activity (*68*). Here, LAG-3 and other co-inhibitory receptors appeared to be expressed to the highest levels on PD1^+^ NK cells, which contained both CD56^++^ and CD56dim cells. PD1^+^ NK cells were the most highly activated NK cells during malaria, and had high granzyme-B and perforin expression, consistent with retained cytolytic capacity. Further studies are required to understand the relationship between CD56neg NK subsets, and PD1^+^ cells. Within γδ T cells, upregulation of co-inhibitory receptors on Vδ2^+^ cells to mediate tolerance has previously been shown in areas of high malaria burden (*11*). Indeed, several of these regulatory genes (*BATF, HAVCR2, TXNIP*) have been previously shown to be increased transcriptionally and at the protein level in γδ T cells in children with recent or high levels of repeated infection (*11, 75, 86*), and here we show that co-inhibitory receptors are also upregulated during an acute infection in a low transmission setting. In areas of high malaria burden, Vδ2^+^ γδ T cells with inhibitory receptors maintain or have enhanced cytolytic capacity and antibody dependent functions (*76*). Consistent with this, co-inhibitory markers were upregulated both transcriptionally and at the protein level was con-current with increased Granzyme-B and perforin, indicative of cytolytic function. Together, these data suggest that regulatory proteins play a role in controlling inflammation, while maintaining other functions of NK and γδ T cells.

Within CD4 and B cell subsets, tolerogenic responses appeared dominated by a major upregulation of IL10. Within CD4 T cells, the Tr1 cell subset was the most highly activated during malaria, and these cells had significantly increased transcription of both canonical cytokines IL10 and IFNγ, and also multiple co-inhibitory receptors (including *LAG3, OX40, TNFR2, GITR, TIM3* and *CTLA4*). A large proportion of malaria-driven DEGs in Tr1 cells were shared with Th1 cells, consistent with an emergence of Tr1 cells from the Th1 cell subset (*101*). Accordingly, DEGs in Tr1 cells also included transcriptional factors with known roles in Tr1 cell development (*MAF, PRDM1 and STING)*(*19, 102–104*). Similarly, within B cell subsets, malaria drove a significant increase in *HIF1A* expression, which has previously been shown to be a critical transcription factor for the induction of IL10 producing Bregs in mouse models (*131*). Consistent with this, we show significantly increased *ex vivo* secretion of IL10 from B cells during malaria compared with 28 days post-treatment. Indeed, during infection B cells were a major contributor of IL10 from lymphocytes during malaria. While IL10 producing CD4 T cells have been well recognised in malaria (*13, 15, 103, 104, 139*), this study is the first to identify B cells as a major source of IL10 during human malaria. Further studies are needed to understand the development of Bregs during malaria, their roles in anti-disease and anti-parasitic (*140*) immunity and/or immunosuppression. The potential link between Tr1 cells as the driver of Breg activation to suppress inflammation and disease as shown in other settings (*141*).

Linking malaria-induced tolerogenic responses across all immune cell subsets, is the importance of Type I IFN signalling, with evidence of upregulation of Type I IFN responses and increased transcription of IFN-stimulated genes across the immune landscape. While first described in viral infection as critical effector cytokines, Type I IFNs also exhibit immunoregulatory effects that impeded control of some non-viral pathogens(*142*), including protozoan parasites (*143*). In malaria infection, Type I IFNs have both protective and detrimental impacts on immune response, parasite clearance and protection from immunopathogenesis, dependent on timing, parasite species and model system (reviewed in (*26*)). However, previous studies in human experimental infection have shown that *P. falciparum* parasites rapidly induce Type I IFNs that enhance development of regulator Tr1 CD4 T cells(*16*). While not directly investigated in malaria, Type I IFN signalling negatively regulates NK cell inflammatory response during viral infection (*71*), promotes upregulation of LAG-3 on NK cells in healthy donors (*68*), and drives Breg induction in helminth infection (*144*). Due to the central role of Type I IFN signalling in driving immunoregulatory responses in malaria, our team is now exploring whether host-directed therapies that transiently block Type I IFNs may have therapeutic roles in enhancing protective anti-parasitic immunity (*145, 146*).

A number of limitations of our study should be noted. Due to low cell numbers contributing to subclustered cell types we were not able to analyse scRNAseq data at the individual cell level, and therefore may have overlooked individual level heterogeneity. To address this, we instead confirmed key transcriptional changes at the protein level in additional patients. The limited number of individuals analysed by scRNAseq also precludes the analysis of the impact of other host factors such as age, sex and/or parasite burden on transcriptional changes. Future studies could take advantage of rapidly developing technologies to increase cell numbers and/or individuals. Additional technical limitations include the use of 3’ sequencing, as such clonal development and TCR/VDJ usage was not investigated. Additionally, only PBMCs responses were assessed, and how these peripheral responses related to the immune response within tissues is unknown. Finally, our study only investigates one study site with all patients >3 years of age and presenting with uncomplicated falciparum malaria, and as such the broad application of results to other malaria transmission settings, young patients and/or disease phenotypes is unknown.

In conclusion, we use scRNAseq analysis of a large number of PBMC cells to make a granular level study of immunoregulatory responses across the immune landscape during falciparum malaria. All data sets and interactive integrated scRNAseq data file is made publicly available for future analysis of the malaria immune landscape by the research community.

## MATERIALS AND METHODS

### Study Design

To investigate malaria driven transcriptional changes in specific immune cell subsets, we sorted live PBMCs from 6 malaria infected donors (day 0), and two subsequent time points after drug treatment (day 7 and 28), along with PBMCs from 2 healthy controls. We performed scRNAseq of these cells, and used clustering and sub-clustering to identify specific immune cell subsets. Differential gene analysis between day 0 and day 28 was performed for each cell cluster and sub-cluster. Key transcriptional changes were confirmed at the protein level with additional donor samples. Patient demographics are in Tables S1, S12 and S13.

### Ethics statement

Ethics approval for the use of human samples was obtained from the QIMR Berghofer Human Research Ethics Committee (HREC P3444), the Northern Territory Department of Health and Menzies School of Health Research ethics committee (HREC 2010-1431), and Medical Research and Ethics Committee, Ministry of Health, Malaysia (NMRR-10-754-6684 and NMRR-12-499-1203). Written informed consent was obtained from all adult study participant or, in the case of children, parents or legal guardians.

### Study participants and peripheral blood mononuclear cell processing

Peripheral blood mononuclear cells (PBMCs) were obtained from patients with acute uncomplicated clinical Plasmodium falciparum malaria from Sabah, Malaysia enrolled in prospective comparative studies of falciparum, vivax and knowlesi malaria, including at three district hospital sites (Kudat, Kota Marudu and Pitas) (147, 148), and a tertiary referral center (Queen Elizabeth Hospital, Kota Kinabalu) (149). This cohort has a high proportion of infected males across all malaria species, possibly because of infection risk of forest worker (147). Patients (aged 3-55) who were positive for Plasmodium species confirmed by microscopy and PCR, who had a fever at the time of presentation or a history of fever in the preceding 24 hours, and who provided consent, were enrolled. Individuals who had been living in the area in the preceding 3 weeks, who were negative for Plasmodium spp. by microscopy and PCR, and who had no history of fever in previous 48 hours were enrolled as endemic healthy controls. Blood was collected in lithium-heparin collection tubes at the time of presentation and follow-up visits at days 7 and 28 after anti-malarial drug treatment (148, 149). PBMCs were isolated from whole blood via density centrifugation with Ficoll-Paque prior to cryopreservation. Samples were selected for this study based on availability.

### 10X Genomics Chromium GEX Library preparation and sequencing

PBMC samples were thawed in RPMI 1640 (Gibco) containing 10% FCS and 0.02% Benzonase. 1E6 PBMCs were stained for viability with Propidium iodide (PI) and live cells were sorted on BD FACSAria™ III Cell Sorter into 2% FBS/PBS and counted on hemocytometer. Up to 10 000 cells were loaded into each lane of Chromium Next GEM Single Cell 3LJ Reagent Kit v3.1 and Gel Bead-in-Emulsion (GEMs) generated in Chromium Controller. Samples were run in two batches, which were later integrated. 3’ Gene Expression Libraries were then generated according to manufacturer’s instructions. Generated libraries were sequenced in a NextSeq 550 System using High Output Kit (150 Cycles) version 1 according to manufacturer’s protocol using paired-end sequencing (150-bp Read 1 and 150 bp Read 2) with the following parameters Read 1: 28 cycles, Index 1: 8 cycles, Read 2: 91 cycles.

### scRNAseq transcriptomic analysis

#### Pre-processing of raw sequencing files

Single cell sequencing data was demultiplexed, aligned and quantified using Cell Ranger version 3.1.0 software (10x Genomics) against the human reference genome (GRCh38-3.0.0), with default parameters. Raw sequencing data and processed Cell Ranger outputs are found at GSE217930, https://www.ncbi.nlm.nih.gov/geo/query/acc.cgi?acc=GSE217930. Cell ranger count matrices for each sample (donor and day) were loaded, merged and analysed using Seurat package v4 (150).Cell cycle scores, mitochondria DNA transcripts and complexity score (log10genes per UMI) were calculated per cell using Seurat built-in functions. Cell cycle score was assigned to each cell using the CellCycleScoring function and evaluated with Principal Component Analysis (PCA). Cells with less than 20% of mitochondria DNA transcripts, complexity score higher than 0.8 and at least 250 genes and 500 UMIs were retained. At the gene level, any hemoglobin-associated genes and genes expressed in less than ten cells were filtered out. The filtered dataset was scaled to regress out the effects of mitochondria DNA transcripts content and cell cycle using the ScaleData function, which regresses each variable individually. We followed the integration workflow included in Seurat to remove unwanted sources of variation. In detail, filtered data was split per donor and day. Each dataset was normalise using the NormalizeData function, and the most variable genes in each of them were selected using FindVariableFeatures. Before integration, the most variable genes shared among the datasets were identified using FindIntegrationAnchors and used to integrate the datasets using IntegrateData function.

#### Cell clustering and sub-clustering

Principal components (PCs) were calculated and the first 30 PCs were used to identify transcriptional clusters and uniform manifold approximation and projection (UMAP) dimensional reduction (‘RunPCA’ and ‘RunUMAP’ functions). Nearest neighbours were calculated and cluster resolution set at 0.6. Cells were annotated based on canonical marker expression, with 15 cell clusters identified. For NK, γδ, CD4 and B cells, sub-clustering analysis was performed. For CD4 T cells, T helper signatures from prior publications; Tfh (89), Tr1 (92) and Th1, Th2, Th17, Treg (90) were analysed within sub clusters. For high level cell clusters, and sub-clustering analysis, marker genes for all clusters (outputs from ‘FindAllMarkers’) are in **Tables S2, S4, S6, S8 and S10**. Processed Seurat file “annotated_Sabah_data_21Oct2022.rds” is found https://doi.org/10.5281/zenodo.6973241

#### Differential gene expression analysis

To identify genes differentially expressed during malaria, ‘FindMarkers’ function with default parameters in Seurat was used comparing specific cell subsets at different time points. DEGs for each cluster and subcluster are found in **Tables S3, S5, S7, S9 and S11**. Identified DEGs were analysed via Gene Set Enrichment Analysis (151) to identify significantly enriched gene ontology (GO) terms for each high-level cluster and subcluster during malaria infection. The HOMER v4.9 package was used to identify significantly overrepresented upstream regulators, of cis-elements 1 kb upstream of the transcription start site (TSS) of the DEGs using the findMotifs.pl script (152).

### Flow cytometric cell phenotyping comparison of scRNAseq samples

2 million cells from the same PBMC vial as used for scRNAseq was used for phenotyping of major cell subsets (**Table S1**). Cells were stained at room temperature with LIVE/DEAD™ Fixable Blue and surface markers with antibodies purchased from BD Biosciences or Biolegend (**Table S12**). Data were acquired with 3-laser Cytek Aurora, and subsets identified as described in **Fig. S1**.

### Flow cytometric ex vivo cytokine production analysis

PBMCs were thawed and 1 million cells incubated at 37°, 5% CO_2_ for 2 hours in additional study patients with falciparum malaria at day 0 and day 28 post-treatment (Table S12). Protein transport inhibitor containing Monensin and protein transport inhibitor containing Brefeldin A were added to cells (both 10 μg/ml, BD Biosciences) and cells cultured for an additional 4 hours. Cell surface staining was performed at RT for 15 minutes with a panel of antibodies purchased from Biolegend, BD Biosciences or Miltenyi (Table S13). Following 2 washes with 2%FCS/PBS, cells were permeabilised with BD cytofix/cytoperm solution for 20 minutes on ice. Intracellular staining (ICS) was performed following this to assess cytokine production. Intracellular staining was performed using antibodies listed in (Table S13). Samples were incubated with the antibodies for 30 minutes on ice, washed twice with BD perm wash buffer then fixed with BD stabilising fixative. All samples were resuspended in 200µl of 2%FCS/PBS to be acquired the following day. Data were acquired using a Cytek Aurora 5, and subsets identified as shown in **Fig. S2 and S8A**. To identify CD16 monocytes, an alternative gating strategy based on CCR2 and CD33 expression (153), due to the rapid down regulation of CD16 in cultured cells.

### Flow cytometric ex vivo cell phenotyping of DEGs

PBMCs were thawed in additional malaria study patients at day 0, and day 28 post malaria infection. Due to sample limitations, not all patients had paired samples for this analysis (**Table S13**). Surface markers, dead cell stains and intracellular stains were performed at the concentrations provided with antibodies purchased from Becton-Dickson Biosciences, Biolegend or Invitrogen (**Table S14**). PBMCs were stained with CD366 (Tim-3) and CD223 (LAG-3) at 37°C for 90 minutes. Cells were then stained at RT for 15 minutes with LIVE/DEAD™ Fixable Blue Dead Cell Stain, washed twice with 2% FCS/PBS and stained at RT for 30 minutes for additional surface markers. Following 2 washes with 2% FCS/PBS, cells were permeabilised with eBioscience™ Fixation/Permeabilization solution for 20 minutes on ice. Intracellular staining (ICS) was performed for 30 minutes on ice following washes to assess intracellular proteases and glycoproteins. Cells were fixed with BD stabilising fixative then resuspended in 200µl of 2%FCS/PBS until acquisition. Data were acquired using a Cytek Aurora 5 and subsets and marker expression identified as described in **Fig. S4A, S4B, S5B, S5C, S7B, and S7C**.

### Flow cytometric analysis

Flow cytometry data were analyzed in FlowJo version 10. Gating strategies are outlined in **Fig. S1, S2, S4A, S5B, S7B-C and S8A**. To measure the co-expression of cytokines or other markers on specific subsets, expression was analysed by SPICE (Simplified presentation of incredibly complex evaluations (46)), and permutation tests between combinations of cytokines/markers performed.

### Statistical analysis

All statistical analysis was performed in RStudio (R version 4.0 or greater). All statistical tests are two-sided. To assess correlations between cellular clusters identified by scRNAseq and flow cytometry, Pearson correlations were calculated. For cell proportions and expression levels, for paired data, Wilcoxon signed-rank test was used and for unpaired data, Mann-Whitney U test was performed.

## Acknowledgments

We thank all the participants and parents of guardians involved in the clinical studies, along with the Malaysian Ministry of Health hospital directors and clinical staff at Kudat, Kota Marudu and Pitas district hospitals and at Queen Elizabeth Hospital, Kota Kinabalu. We thank support staff in QIMR Flow Cytometry and Imaging Facility, QIMR Sample Processing and Sequencing Service, and Dr. Jessica Engel for laboratory support.

## Author contributions

Conceptualization: TGC, JRL, MJB

Methodology: TGC, JRL, MJB

Software: NLD, TGC, ZP, MJB

Validation: NLD, TGC, ZP, JRL, JH, DA, MSFS, MJB.

Formal analysis: NLD, TGC, ZP, JRL, DA, JH, MJB

Investigation: NLD, TGC, ZP, KB, JRL, DA, MSFS, AS, MJB

Resources: KP, TW, BB, MJG, NA,

Data Curation : NLD, TGC, ZP, MJB

Writing - Original Draft: NLD, TGC, MJB

Writing - Review & Editing: NLD, JRL, MS, BB, JAL, CRE, NMA, MJB

Visualization: NLD, TGC, JRL, JH, MJB

Supervision: AL, CE, MJB

Project administration : MJB,

Funding acquisition: JRL, NMA, MJB

All authors have read and approved the final version of the manuscript. MJB approves this version of the manuscript on behalf of TGC (deceased).

## Data and materials availability

Raw sequencing data and processed Cell Ranger outputs are found GSE217930, https://www.ncbi.nlm.nih.gov/geo/query/acc.cgi?acc=GSE217930

Processed Seurat file “annotated_Sabah_data_21Oct2022.rds” is found https://doi.org/10.5281/zenodo.6973241

## Supplementary Materials

Fig. S1. Gating strategy of flow cytometric cell phenotyping comparison of scRNAseq samples.

Fig. S2. Innate cell subset gating strategy for *ex vivo* cytokine analysis.

Fig. S3. GSEA of DEGs from NK cell subsets.

Fig. S4. Flow cytometry analysis of NK cells.

Fig. S5. Flow cytometry analysis of γδ T cells.

Fig. S6. Proportional distribution of CD4 T cell subsets.

Fig. S7. Proportional distribution of B cell subsets and CD98 protein expression.

Fig. S8. Intracellular cytokine expression by major lymphocyte subsets.

Table S1. Patient characteristics for scRNAseq.

Table S2. PBMC cluster marker genes (attached file)

Table S3. PBMC cluster DEGs (attached file)

Table S4. NK subset marker genes (attached file)

Table S5. NK subset DEGs (attached file)

Table S6. γδ T cell subset marker genes (attached file)

Table S7. γδ T cell subset DEGs (attached file)

Table S8. CD4 T cell subset marker genes (attached file)

Table S9. CD4 T cell subset DEGs (attached file)

Table S10. B cell subset marker genes (attached file)

Table S11. B cell subset DEGs (attached file)

Table S12. Patient characteristics for *ex vivo* cytokine production analysis.

Table S13. Patient characteristics for *ex vivo* cell phenotyping

Table S14: Antibodies for *ex vivo* cell phenotyping comparison of scRNAseq samples

Table S15: Antibodies for *ex vivo* cytokine analysis

Table S16: Antibodies for *ex vivo* cell phenotyping analysis

