## Supplementary Figures and Tables for "Malaria drives unique regulatory responses across multiple immune cell subsets"

**Supplementary Materials**

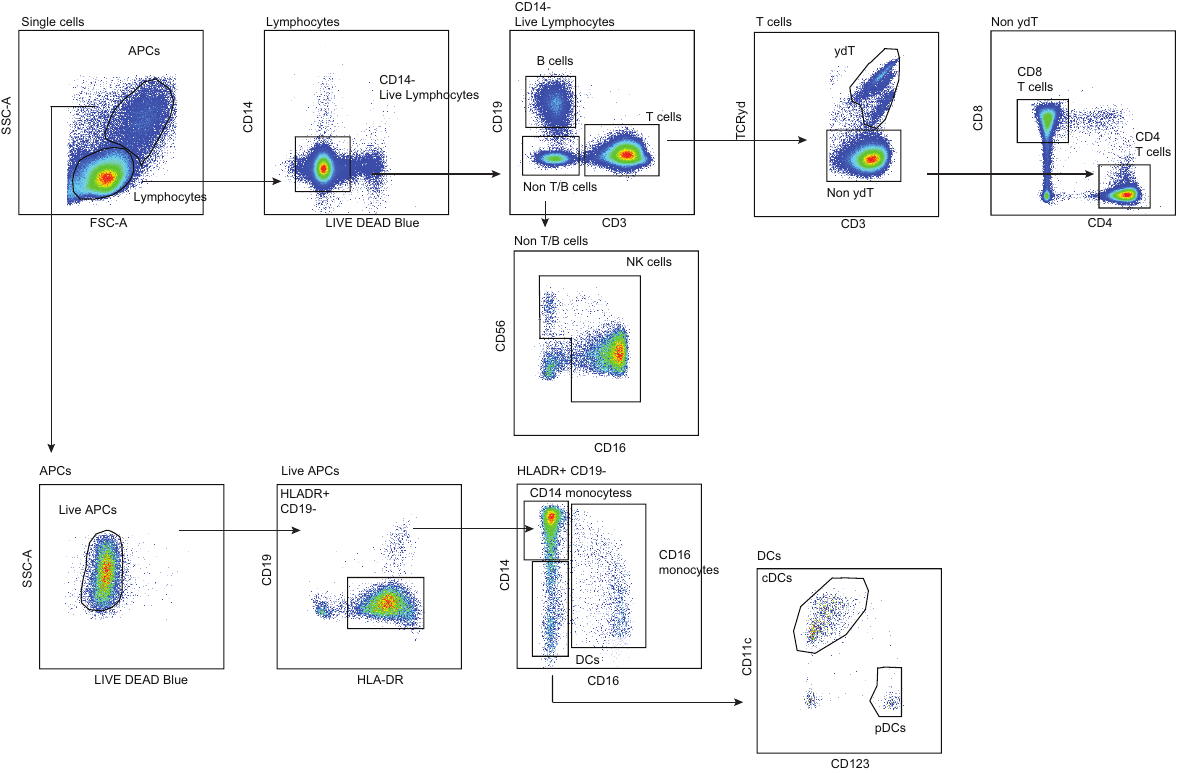

**Fig. S1. Gating strategy of Flow cytometric cell phenotyping comparison of scRNAseq samples.** PBMCs from each individual analysed for scRNAseq were analysed by flow cytometry to identify major cell subsets. Gating strategy is outlined.

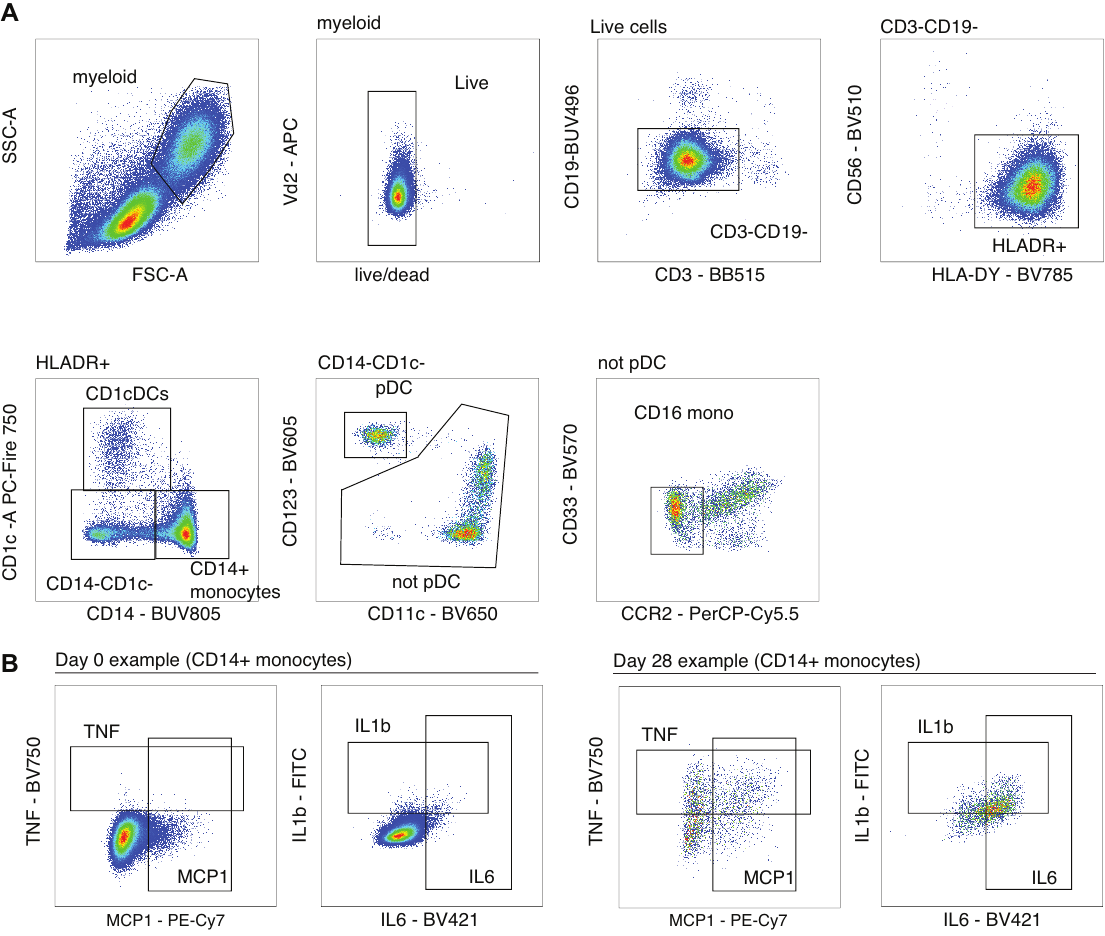

**Fig. S2. Innate cell subset gating strategy for ex vivo cytokine analysis.** To investigate key changes to innate myeloid cells at the protein level PBMCs from patients with malaria (day 0) and 28 days post treatment where cultured ex vivo for 4 hours with monensin, and cytokine production measured by flow cytometry. (**A**) Gating strategy to identify CD14^+^ and CD16^+^ monocytes, and CD1c^+^ DCs. Due to the down regulation of CD16 in culture cells, alternative gating strategy as HLA-DR^+^/CD1c^-^/CD14^-^/CD123^-^/CCR2^-^/CD33^-^ was used. (**B**) TNF, MCP1, IL6, IL1β expression from CD14 monocytes at day 0 and day 28.

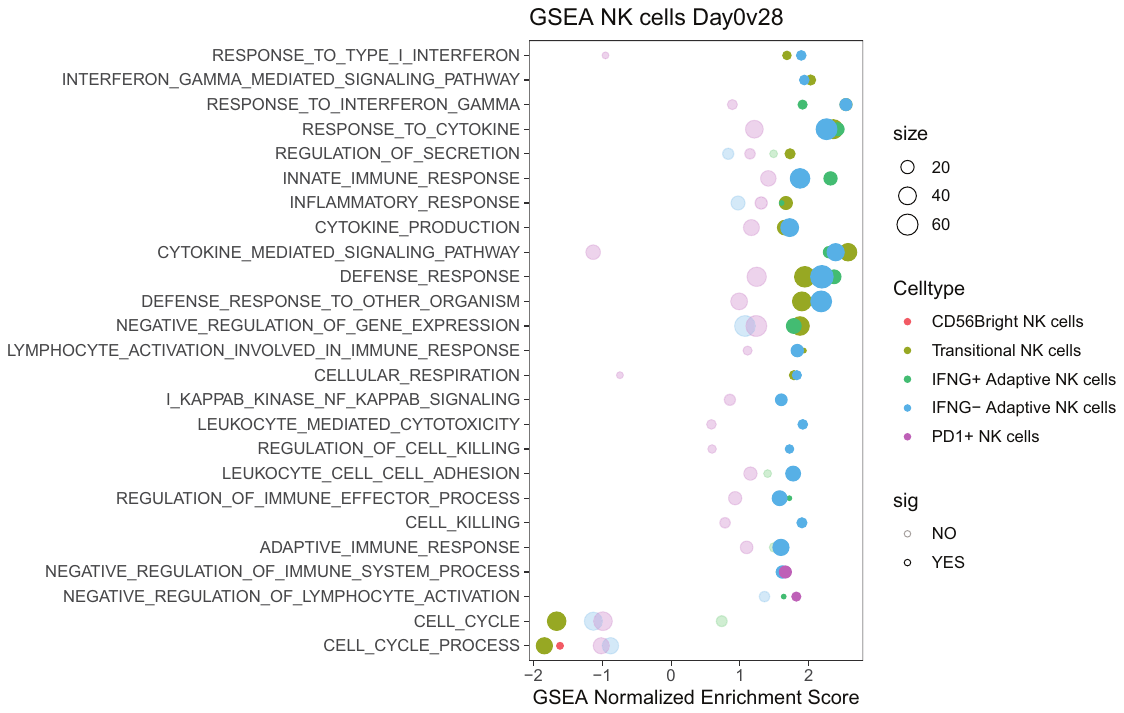

**Fig. S3. GSEA of DEGs from NK cell subsets**. DEGs identified between day 0 and day 28 for each NK cell subset were analysed by Gene Set Enrichment Analysis. Enriched signatures for each subset indicated.

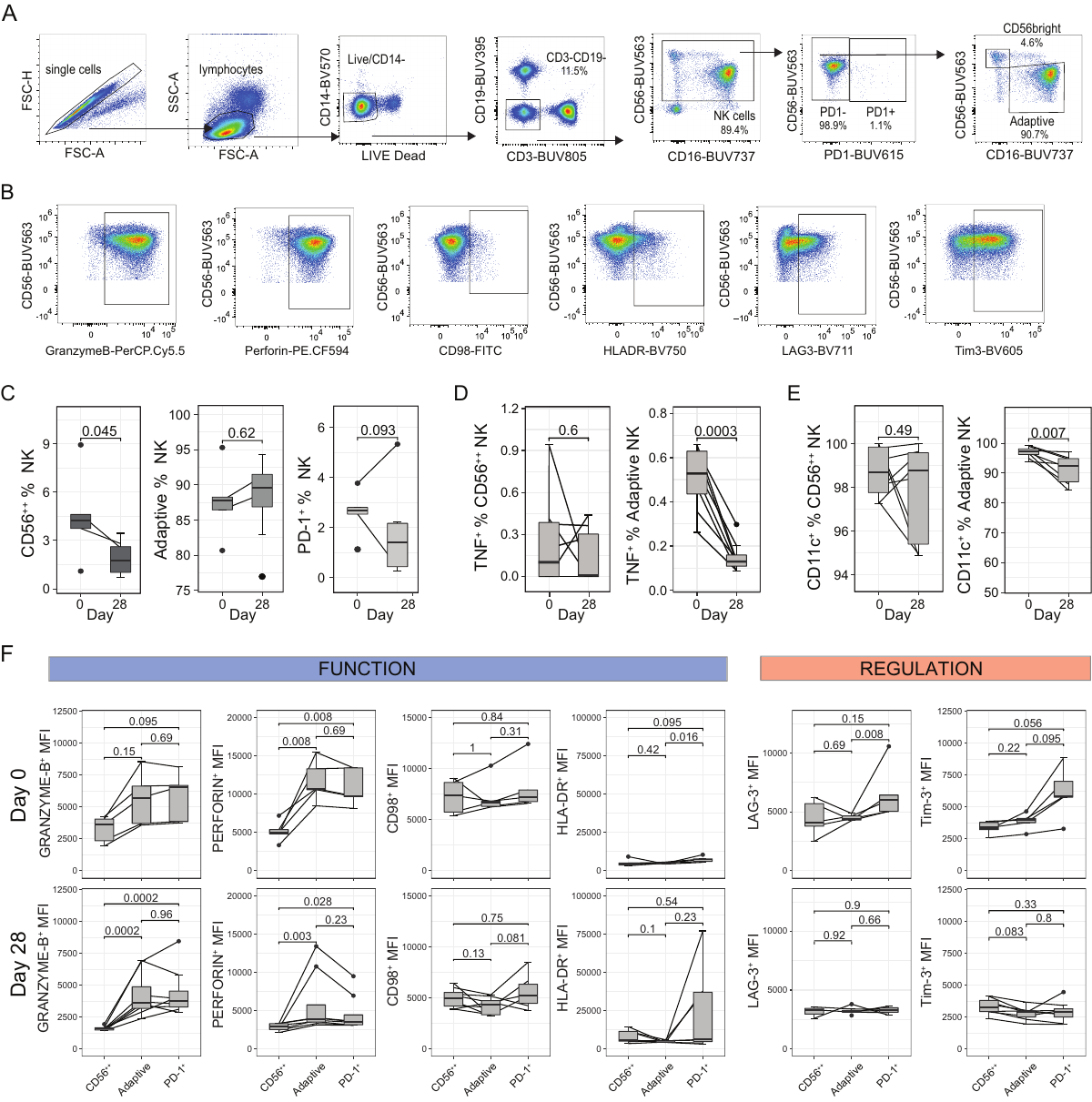

**Fig. S4. Flow cytometry analysis of NK cells.** Peripheral blood mononuclear cells from individuals with acute P. falciparum infection (Day 0, n = 5), 28-days post-infection (Day 28, n = 8) were analysed ex vivo to identify NK cell protein expression of DEGs. (**A**) Gating strategy to identify PD1^+^, CD56 bright (CD56^++^) and Adaptive NK subsets. (**B**) Levels of expression of Granzyme-B, Perforin, CD98, ICOS, HLA-DR, LAG-3 and Tim-3 against CD56 expression at day 0 in total NK (expressed by MFI). (**C**) Proportion of NK subsets within the total NK cell population at day 0 and day 28. (D) TNF expression on CD56^++^ and Adaptive NK cells at day 0 and day 28. (**E**) CD11c expression on CD56bright and Adaptive NK cells at day 0 and day 28. (**F**) Levels of expression (expressed as MFI) of Granzyme-B, Perforin, CD98, HLA-DR, LAG-3 and Tim-3 on CD56^++^, Adaptive and PD1^+^ NK cells at day 0 and day 28. For all data, box plots show the median and IQR of volunteers, lines represent paired observations, group comparisons performed by Mann Whitney U test and paired group comparisons performed by Wilcoxon signed rank test.

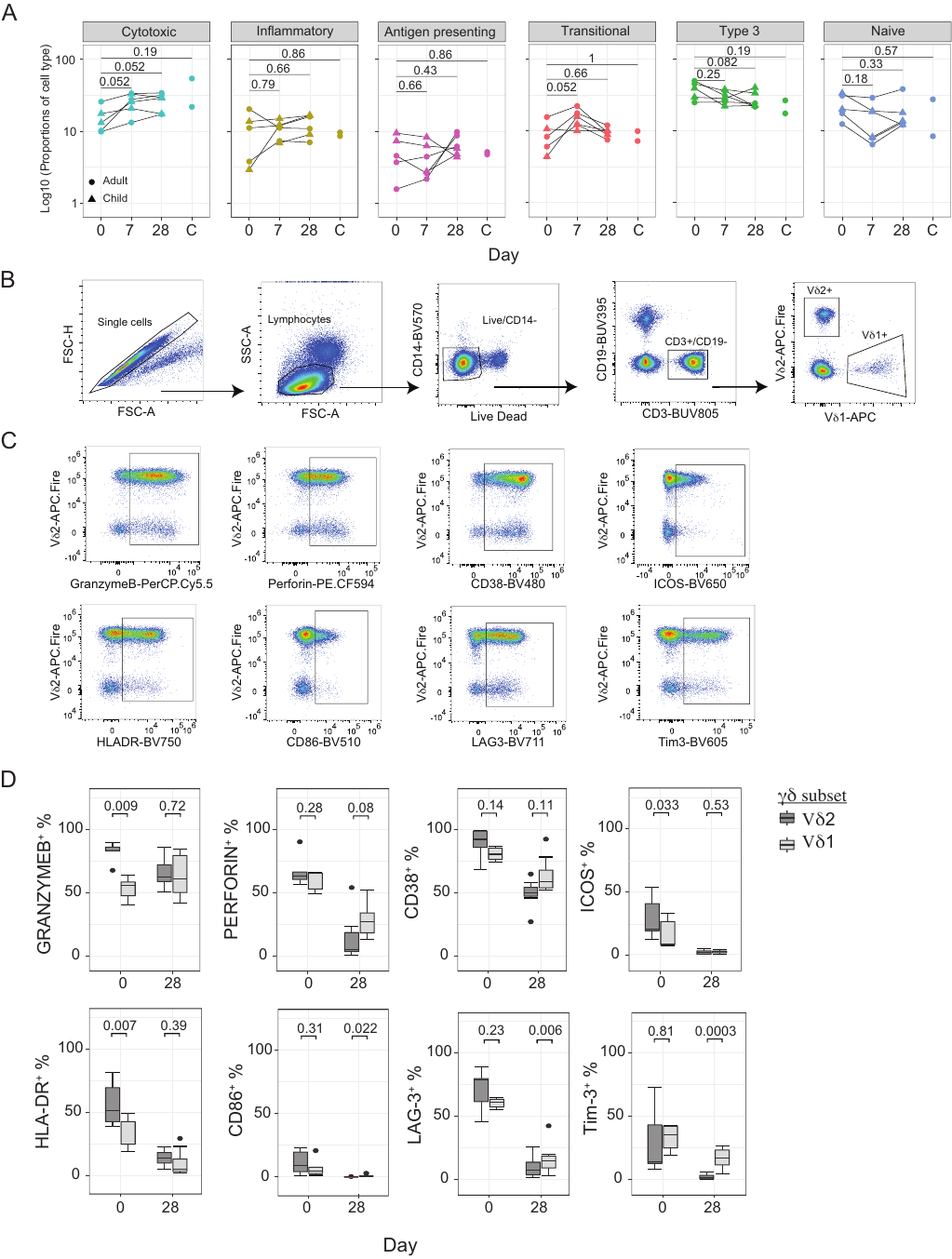

**Fig. S5. Flow cytometry analysis of γδ T cells.** (**A**) Relative proportions of identified of γδ T cells subsets within scRNAseq data during malaria infection (day 0) and day 7, and 28 day post treatment, and in healthy uninfected individuals. (**B**) Gating strategy to identify Vδ2 and Vδ1 γδ T cells, and measure marker expression. (**C**) Levels of expression of Granzyme-B, Perforin, CD38, ICOS, HLA-DR, CD86, LAG-3, Tim-3 between Vδ2 and Vδ1 γδ T cells at day 0 and day 28 (expressed by MFI). D) Surface marker frequencies comparison between Vδ2 and Vδ1 γδ T cells at day 0 and day 28 timepoints. For all data, box plots show the median and IQR of volunteers, lines represent paired observations, group comparisons performed by Wilcoxon rank sum test.

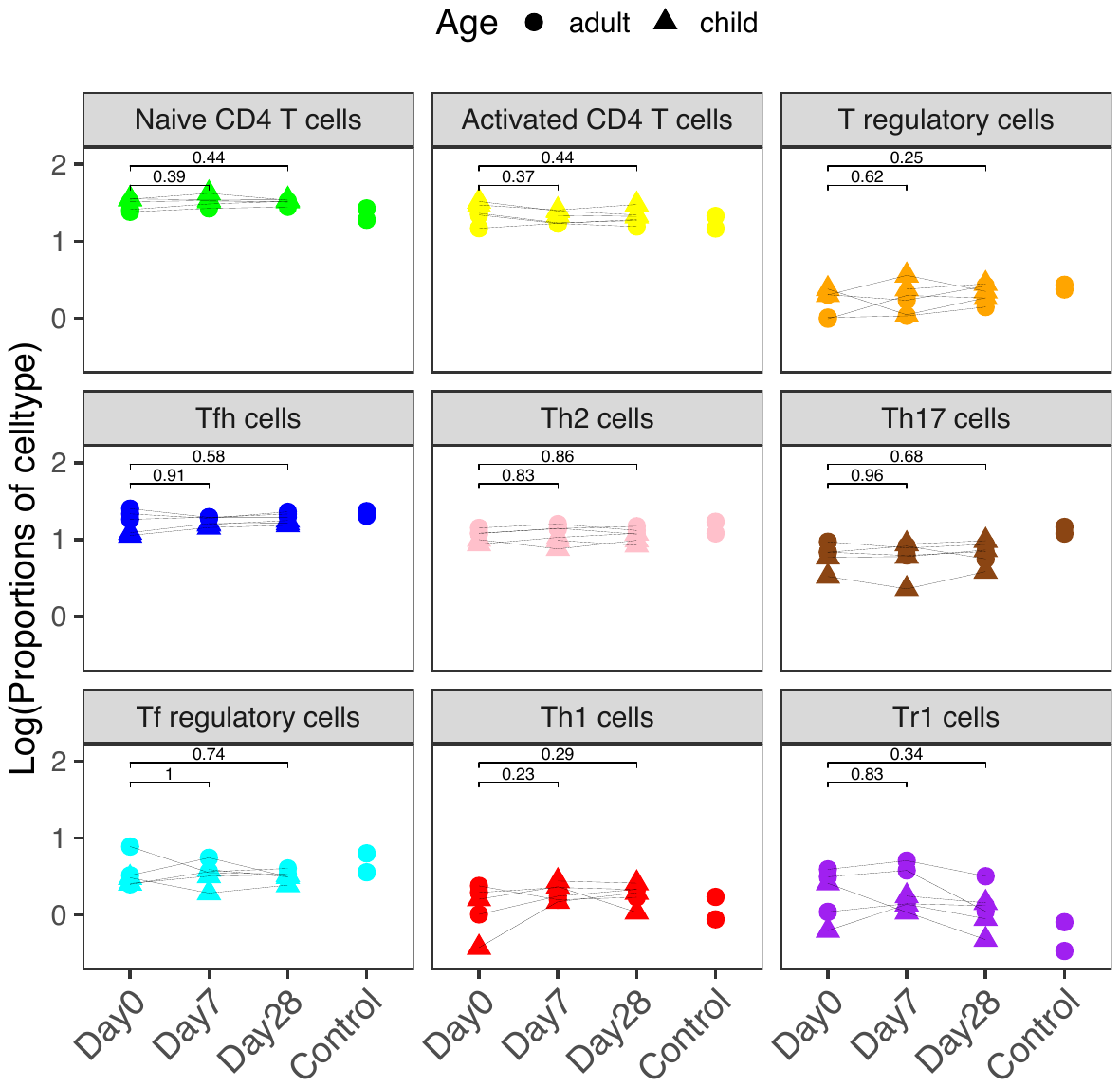
**Fig. S6. Proportional distribution of CD4 T cell subsets.** The proportion of each CD4 T cell subset, as a percentage of total CD4 T cells was calculated for each individual and timepoint. Log10 proportion is show. Group comparisons were performed by wilcoxon rank sum test.

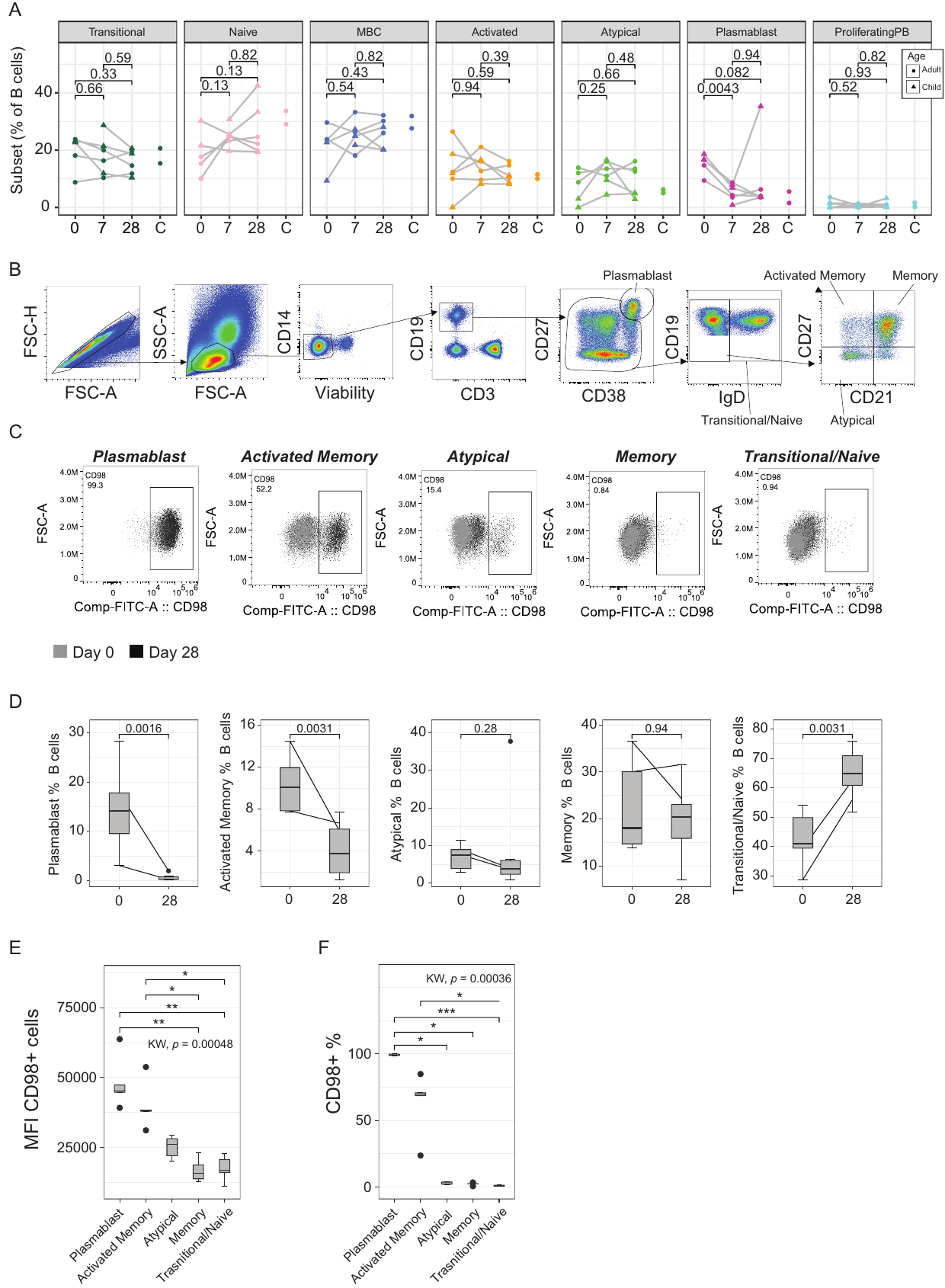

**Fig. S7. Proportional distribution of B cell subsets and CD98 protein expression.** (**A**) The proportion of each B cell subset, as a % of total B cells a was calculated for each individual and timepoint. Log10 proportion is show. P is Wilcox rank sum test. (**B**) Gating strategy for identifying B cell subsets, Concatenated sample example. (**C**) CD98 expression on each B cell subset at day 0 and day 28, concatenated samples. (**D**) The proportion of each B cell subset, as a % of total B cells at day 0 (n=5) and day 28 (n=8). (**E**) MFI of CD98^+^ B cell subset and (**F**) CD98 % B cell subset expression at day 0. Kruskal-Wallis and post-hoc Dunn test (FDR adjusted) indicated.
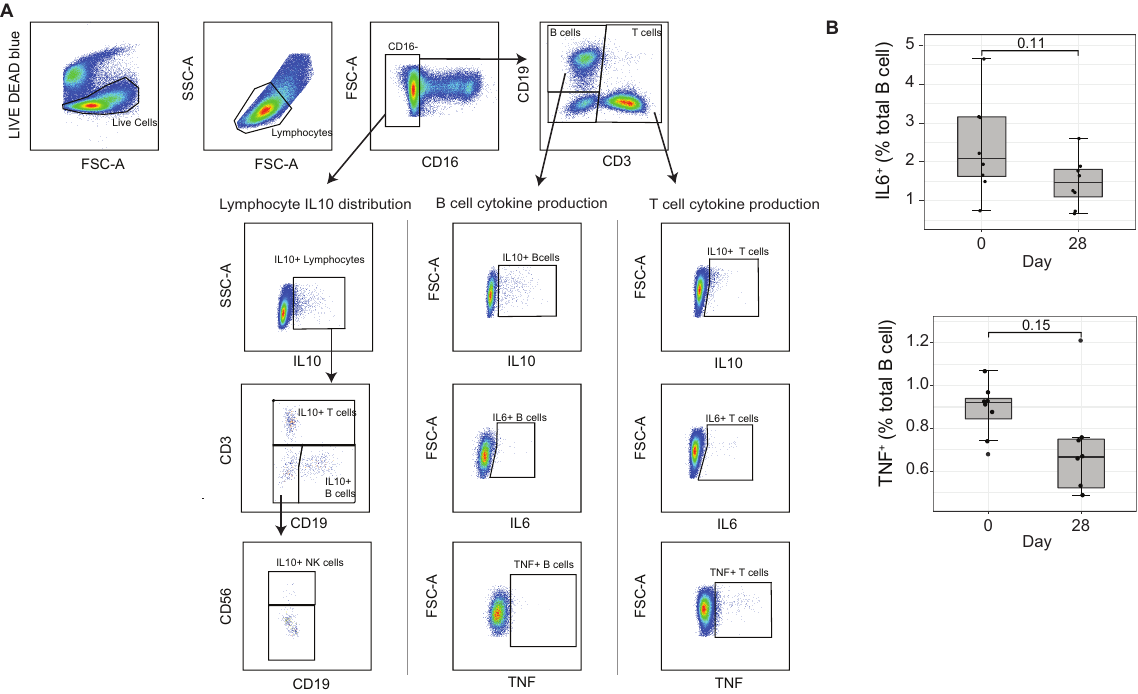

**Fig. S8. Intracellular cytokine expression by major lymphocyte subsets.** (**A**) Gating strategy and representative examples of IL10, IL6 and TNF intracellular cytokine expression across major lymphocyte subsets. (**B**) IL6 and TNF expression in B cells at day 0 (n=8) and day 28 (n=8). Wilcoxon signed rank test indicated.

|  |  |  |  |  | **Cells passed QC** | | |
| --- | --- | --- | --- | --- | --- | --- | --- |
| **10X ID** | **Sample** | **Age** | **Sex** | **Parasitemia** | **Day 0** | **Day 7** | **Day 28/**  **control** |
| E/F/G | Child 1 | 6 | M | 110730 | 0 | 2553 | 6961 |
| M/S/N | Child 2 | 8 | M | 64000 | 6274 | 7726 | 5833 |
| O/T/P | Child 3 | 13 | M | 770 | 2364 | 5510 | 5209 |
| A/B/C | Adult 1 | 21 | M | 26076 | 2746 | 5435 | 5574 |
| I/Q/J | Adult 2 | 22 | F | 23603 | 2938 | 7082 | 4389 |
| K/R/L | Adult 3 | 24 | M | 34 | 1870 | 6513 | 6320 |
| D | Control 1 | 27 | M | - |  |  | 11752 |
| H | Control 2 | 20 | M | - |  |  | 9027 |

**Table S1: Patient characteristics for scRNAseq.**

**Table S2: PBMC cluster marker genes (attached file)**

**Table S3: PBMC cluster DEGs (attached file)**

**Table S4: NK subset marker genes (attached file)**

**Table S5: NK subset DEGs (attached file)**

**Table S6: γδ T cell subset marker genes (attached file)**

**Table S7: γδ T cell subset DEGs (attached file)**

**Table S8: CD4 T cell subset marker genes (attached file)**

**Table S9: CD4 T cell subset DEGs (attached file)**

**Table S10: B cell subset marker genes (attached file)**

**Table S11: B cell subset DEGs (attached file)**

| **Patient** | **Age** | **Sex** | **Parasitemia** |
| --- | --- | --- | --- |
| 1 | 28 | M | 17208 |
| 2 | 38 | F | 6254 |
| 3 | 37 | M | 1862 |
| 4 | 30 | M | 56922 |
| 5 | 23 | M | 23868 |
| 6 | 54 | F | 3317 |
| 7 | 53 | M | 7563 |
| 8 | 30 | M | 5600 |
| Population | 33.5  (29.5-41.75)  median, IQR | 62.5% Male | 6908.5  (5029.25- 18873)  Median, IQR |

**Table S12: Patient characteristics for *ex vivo* cytokine production analysis.**

| **Patient** | **Age** | **Sex** | **Parasitemia** | **Day** |
| --- | --- | --- | --- | --- |
| 9 | 45 | M | 81062 | 0 |
| 10 | 37 | F | 32267 | 0 |
| 11 | 39 | M | 1508 | 0 |
| 12 | 54 | F | 27905 | 0/28 |
| 13 | 36 | F | 17263 | 0/28 |
| 14 | 49 | M | 17759 | 28 |
| 15 | 54 | M | 84403 | 28 |
| 16 | 39 | M | 40895 | 28 |
| 17 | 46 | F | 45844 | 28 |
| 18 | 16 | M | 1464 | 28 |
| 19 | 21 | F | 9648 | 28 |
| Population | 39  (36.5-47.5)  median, IQR | 54.5% Male | 27905  (13455-43370)  Median, IQR |  |

**Table S13: Patient characteristics for *ex vivo* cell phenotyping**

| **Target** | **Fluorochrome** | **Clone** | **Manufacturer** | **Catalogue number** | **Dilution** |
| --- | --- | --- | --- | --- | --- |
| CD16 | BUV395 | 3G8 | BD | 563785 | 1/500 |
| CD45RA | BUV563 | HI100 | BD | 565702 | 1/1000 |
| CD4 | BUV737 | SK3 | BD | 612748 | 1/250 |
| CD27 | BV421 | M-T271 | BD | 562513 | 1/250 |
| CD123 | PacBlue | 6H6 | Biolegend | 306043 | 1/50 |
| CD56 | BV510 | HCD56 | Biolegend | 318340 | 1/50 |
| CD127 | BV570 | A019D5 | Biolegend | 351307 | 1/80 |
| CD11c | BV650 | Bu15 | Biolegend | 337238 | 1/500 |
| CD8 | BV711 | RPA-T8 | BD | 563677 | 1/1000 |
| CD19 | BV750 | HIB19 | Biolegend | 302261 | 1/500 |
| HLA-DR | BV785 | L243 | Biolegend | 307642 | 1/250 |
| TCR γδ | FITC | B1 | Biolegend | 331208 | 1/50 |
| CD3 | AF532 | UCHT1 | Invitrogen | 58-0038-42 | 1/50 |
| CD14 | PerCP-Cy5.5 | M5E2 | Biolegend | 301824 | 1/100 |
| ICOS | PE | DX29 | BD | 557802 | 1/100 |
| CD86 | PE-Daz594 | IT2.2 | Biolegend | 305434 | 1/250 |
| CXCR5 | PE-Cy7 | J252D4 | Biolegend | 356924 | 1/250 |
| CD38 | AF647 | HIT2 | Biolegend | 303514 | 1/500 |
| CD25 | APC-R700 | 2A3 | BD | 565106 | 1/250 |
| Vδ2 | APC/Fire 750 | B6 | Biolegend | 331420 | 1/500 |

**Table S14: Antibodies for *ex vivo* cell phenotyping comparison of scRNAseq samples**

| **Antibody** | **Fluorochrome** | **Clone** | **Manufacturer** | **Catalogue number** | **Dilution** |
| --- | --- | --- | --- | --- | --- |
| Surface staining | | | | | |
| CD19 | BUV496 | SJ25C1 | BD | 612938 | 1/100 |
| CD64 | BUV737 | 10.1 | BD | 564425 | 1/50 |
| CD14 | BUV805 | M5E2 | BD | 612902 | 1/50 |
| CD86 | BV480 | 2331 | BD | 566131 | 1/50 |
| CD3 | BB515 | UCHT1 | BD | 564466 | 1/100 |
| CD56 | BV510 | HCD56 | Biolegend | 318340 | 1/50 |
| CD33 | BV570 | WM53 | Biolegend | 303417 | 1/50 |
| CD123 | BV605 | 6H6 | Biolegend | 306026 | 1/100 |
| CD11c | BV650 | B915 | Biolegend | 117310 | 1/200 |
| HLA-DR | BV785 | L243 | Biolegend | 307642 | 1/100 |
| CCR2 | PerCPCy5.5 | K036C2 | Biolegend | 357204 | 1/50 |
| Vδ2 | APC | B6 | Biolegend | 331418 | 1/50 |
| CD16 | AF700 | 3C78 | Biolegend | 302026 | 1/100 |
| CD1c | APC-FIRE | L161 | Biolegend | 331545 | 1/30 |
| Live/Dead | - | - | Invitrogen | L23105 | 1/5000 |
| Intracellular staining | | | | | |
| IFN-γ | BUV395 | B27 | BD | 563563 | 1/25 |
| IL-6 | BV421 | C8-6 | BD | 501119 | 1/100 |
| IL-4 | BV711 | MD4–25D2 | BD | 564112 | 1/25 |
| TNF | BV750 | Mab11 | BD | 566359 | 1/100 |
| IL-12 | AF647 | 503-F7 | BD | 565023 | 1/25 |
| IL-10 | PE-Dazzle | JES–19F1 | BD | 506812 | 1/25 |
| MCP-1 | PE-Cy7 | 5D3-F7 | Biolegend | 502614 | 1/100 |
| IFN-α | PE | LT27-295 | Miltenyi | 130-092-602 | 1/100 |
| IL-1β | FITC | CRM56 | Invitrogen | 11-7018-42 | 1/50 |

**Table S15: Antibodies for *ex vivo* cytokine analysis**

| **Antibody** | **Fluorochrome** | **Clone** | **Manufacturer** | **Catalogue number** | **Dilution** |
| --- | --- | --- | --- | --- | --- |
| Surface staining | | | | | |
| CD366 (Tim-3) | BV605 | F38-2E2 | Biolegend | 345017 | 1 in 25 |
| CD223 (LAG-3) | BV711 | 11C3C65 | Biolegend | 369319 | 1 in 50 |
| CD27 | AF700 | M-T271 | BD | 560611 | 1 in 100 |
| Vδ1 | APC | TS8.2 | Invitrogen | 17-5679-42 | 1 in 200 |
| Vδ2 | APC-Fire | B6 | Biolegend | 331420 | 1 in 100 |
| CD19 | BUV395 | SJ25C1 | BD | 563549 | 1 in 100 |
| CD56 | BUV563 | NCAM16.2 | BD | 612928 | 1 in 500 |
| CD279 (PD1) | BUV615 | EH12.1 | BD | 612991 | 1 in 50 |
| CD16 | BUV737 | 3G8 | BD | 612787 | 1 in 500 |
| CD3 | BUV805 | SK7 | BD | 612893 | 1 in 200 |
| CD57 | BV421 | NK-1 | BD | 563896 | 1 in 500 |
| CD38 | BV480 | HIT2 | BD | 566137 | 1 in 500 |
| CD86 | BV510 | 2331 (FUN-1) | BD | 563461 | 1 in 100 |
| CD14 | BV570 | M5E2 | Biolegend | 301831 | 1 in 50 |
| CD278 (ICOS) | BV650 | DX29 | BD | 563832 | 1 in 50 |
| HLA-DR | BV750 | L243 | Biolegend | 307672 | 1 in 200 |
| IgD | BV785 | IA6-2 | Biolegend | 348242 | 1 in 200 |
| CD98 | FITC | MEM-108 | Biolegend | 315603 | 1 in 50 |
| LIVE/DEAD Fixable Blue Dead Cell Stain | - | - | Invitrogen | L23105 | 1 in 2500 |
| Intracellular staining | | | | | |
| Perforin | PE-CF594 | δG9 | BD | 563763 | 1 in 5000 |
| Granzyme-B | PerCP-Cy5.5 | QA16A02 | Biolegend | 372212 | 1 in 500 |

**Table S16: Antibodies for *ex vivo* cell phenotyping analysis**
